# Origins and diversity of pan-isotype human bone marrow plasma cells

**DOI:** 10.1101/2024.05.08.592267

**Authors:** Gaspar A. Pacheco, Vishal Rao, Duck Kyun Yoo, Shahab Saghaei, Pei Tong, Sachin Kumar, Orlee Marini-Rapoport, Zahra Allahyari, Ali S. Moghaddam, Romina Esbati, Aida Alirezaee, Aric Parnes, Sarita U. Patil, Duane R. Wesemann

## Abstract

Bone marrow plasma cells (BMPCs) produce durable, protective IgM, IgG, and IgA antibodies, and in some cases, pro-allergic IgE antibodies, but their properties and sources are unclear. We charted single BMPC transcriptional and clonal heterogeneity in food-allergic and non-allergic individuals across CD19 protein expression given its inverse correlation to BMPC longevity. Transcriptional and clonal diversity revealed distinct functional profiles. Additionally, distribution of somatic hypermutation and intraclonal antibody sequence variance suggest that CD19low and CD19high BMPCs arise from recalled memory and germinal center B cells, respectively. Most IgE BMPCs were from peanut-allergic individuals; two out of 32 from independent donors bound peanut antigens in vitro and in vivo. These findings shed light on BMPC origins and highlight the bone marrow as a source of pathogenic IgE in peanut allergy.

## Introduction

Antibodies circulating in the bloodstream are indicative of learned immunity from prior infection, vaccination, and in some cases, exposure to allergens. Steady state levels of antibodies over time can play a major role in influencing susceptibility to infection (*1, 2*) and allergies (*3*). Bone marrow plasma cells (BMPCs) are a long-lived source of stable circulating antibodies, with longer-lasting subsets having undergone germinal center (GC) affinity maturation. Longer-lasting subsets exhibit low CD19 expression (*4*), which is thought to arise from CD19-expressing BMPCs progressively losing CD19 from the plasma cell surface (*5*). While it is known that recalled immunity can lead to more durable long-lived antibody levels (*6, 7*), a mechanistic link between recalled immunity and greater levels of CD19^low^ plasma cells is unclear.

In addition, while BMPC production of IgM, IgG, and IgA plays an important role in steady-state protection from infectious invasion, long-lived production of IgE in allergic individuals may sensitize them to dangerous allergic attacks. Of particular intrigue is the unique regulation of IgE production in comparison to production of other antibody isotypes. Upon class switch recombination to IgE, B cells rapidly differentiate to short-lived plasma cells before quickly vanishing (*8–10*). By contrast, IgG B cells can maintain a stable memory pool in circulation and give rise to long-lived plasma cells. Maintenance of allergic memory can be possible through secondary heavy chain class switch recombination from allergen-specific IgG memory cells (*10–14*). IgE BMPCs have been detected in allergic individuals (*15, 16*). However, cells producing food allergen-specific IgE have only been detected in peripheral blood and intestinal mucosa to date (*17–20*), where B cell activation and IgH class-switching can occur. Given that many individuals have severe and persistent allergy across decades, a bone marrow source for pathogenic, IgE-producing cells has long been speculated (*15, 16, 21–23*). While T cell help is associated with long-lived BMPC generation and is likely necessary for IgE production (*24–26*) in humans, the contribution of BMPCs to pathogenic IgE in food allergy remains unknown (*27*). Our analysis of allergic and non-allergic bone marrow samples here sheds new light on the likely sources of long-lived plasma cells and identifies the bone marrow as a source of pathogenic IgE antibodies in peanut allergy.

### Human BMPCs exhibit a heterogeneous transcriptional landscape

To study the source dynamics of bone marrow plasma cells and to gain insights into potential long-lived IgE production, we collected bone marrow samples from nine individuals, six with food allergy and three without any history of food allergy (Supplementary Table 1). We performed VDJ and 5’ gene expression single-cell RNA sequencing paired with CITE-seq on CD138-enriched, FACS-purified BMPCs (Fig. 1A & S1). We sequenced 60,548 individual BMPCs and identified 11 Ig gene-independent BMPC clusters with comparable numbers of genes and transcripts (unique molecular identifiers, UMIs) expressed in each cluster, as well as a low proportion of mitochondrial gene expression and detectable Ig expression (Fig. 1B-C & S2). Cells showed robust expression of plasma cell-defining transcription factors and markers (*28–31*) and lacked expression of B cell lineage-defining genes, confirming their plasma cell identity (Fig. 1D-F). All clusters contained BMPCs expressing different heavy chain (IgH) and light chain (IgL) isotypes but differed in IgH somatic hypermutation (SHM) (Figs. 1G-I, S2, & S3) and surface CD19 protein levels (Fig. 1J). We identified 54 IgE BMPCs which were primarily found in cluster 6 and to a lesser extent in cluster 8 (Fig. 1G). Analysis of B cell receptor (BCR) sequences identified around half of all BMPC sequences as singlets. Within expanded clonal families, we identified a high level of connectivity of shared clonotypes between clusters, as previously found for mouse BM LLPCs (*32*) (Fig. 1K) and moderate connectivity between different IgH isotypes, of which IgA1, IgG1, and IgG2 were the most common among clonal families that spanned multiple isotypes (Fig. S3). The 20 largest clonal families (clone size 24-132 cells) spanned multiple clusters and were dominated by IgA1 or IgG1 BMPCs, with relatively minor contributions of other isotypes (Fig. S3).

**Fig. 1.**
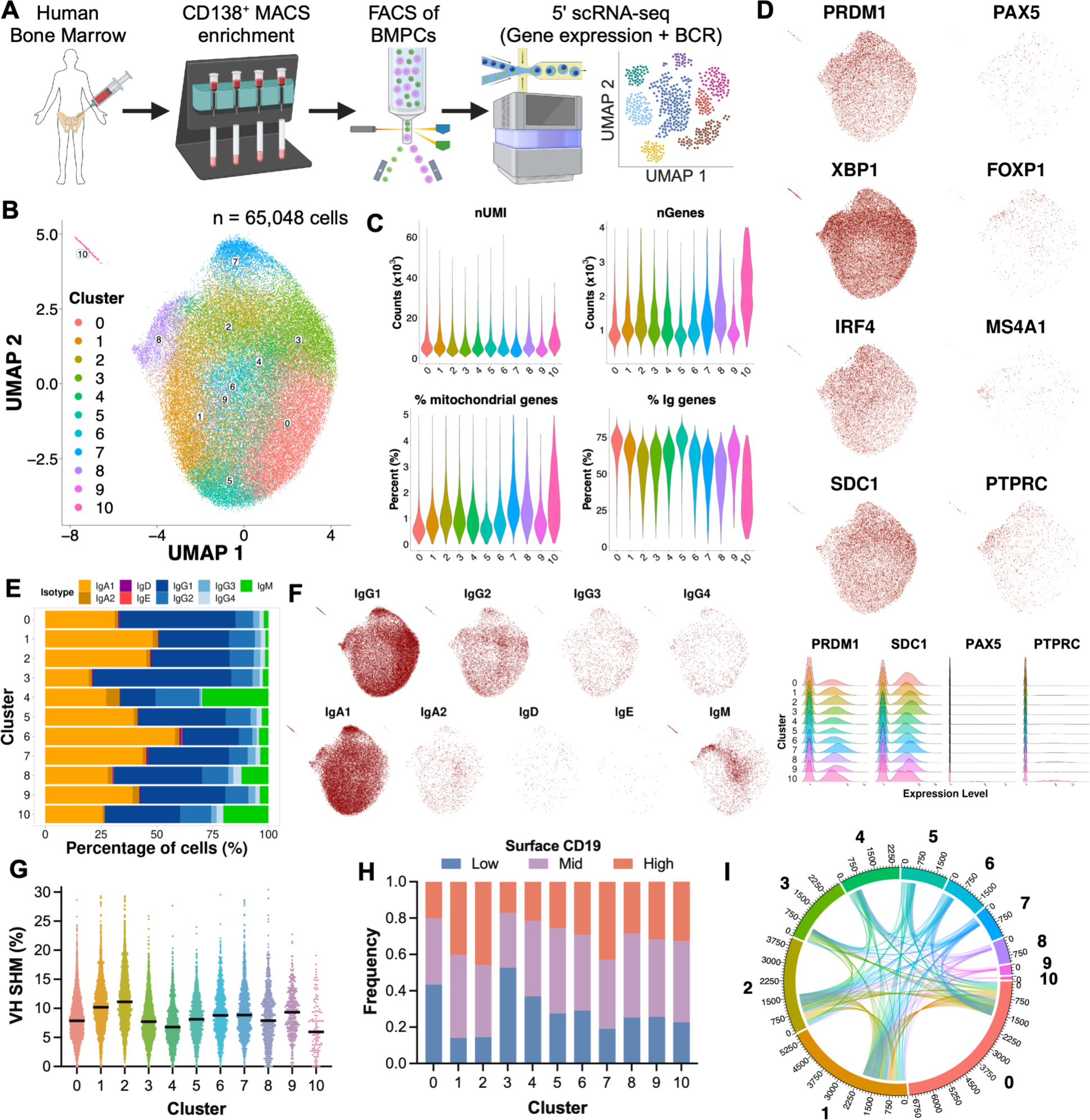
scRNA-seq reveals intrinsic human bone marrow plasma cell heterogeneity. (**A**) Study overview. (**B**) UMAP projection of sorted BMPCs, colored by cluster assignment (n = 65,048 cells). Clustering was performed after exclusion of immunoglobulin genes. (**C**) Quality control of clustered cells (number of RNA molecules and total genes detected, and percentage of mitochondrial and immunoglobulin genes). (**D-F**). Expression of plasma cell- and B cell-associated genes among clusters. (**G-H**) IgH isotypic composition of clustered cells. (**I**) IgH somatic hypermutation among clusters. (**J**) Frequency of binned surface CD19 protein expression of BMPC clusters. (**K**) Clonal relationships between clusters. Polar axis represents number of cells with V(D)J arrangement available in each cluster.

We identified differentially expressed genes (DEGs) for each cluster (Fig. 2A-B). Some clusters distinctly expressed cellular features associated with metabolism (cluster 6), MHC-II expression (cluster 8), interferon response (cluster 9), and proliferation (cluster 10). Proliferating cells in cluster 10 likely represent plasmablasts. We further investigated cluster 6 given its association with IgE and what appeared to be a tendency toward high metabolic demand. Gene set enrichment analysis (GSEA) identified amino acid biosynthesis, ER stress responses, and translation as key upregulated processes, while genes associated with cell migration were underrepresented in cluster 6 compared to all other BMPCs (Fig. 2C-D). Indeed, genes associated with amino acid transport, transamination, folate synthesis, and mitochondrial unfolded protein response (UPR) were more highly expressed in cluster 6 (Fig. 2E), and amino acid availability has been identified as a limiting step in ASC generation (*33*). Similarly, genes encoding translation factors were highly expressed in cluster 6 (Fig. 2F), while molecules associated with cell migration and adhesion in plasma cells were downregulated (Fig. 2G) (*34*). Interestingly, cluster 4 highly expressed *CCR10* and had the highest proportion of IgM and IgA2 plasma cells, which is consistent with a gut origin (*35*).

**Fig. 2.**
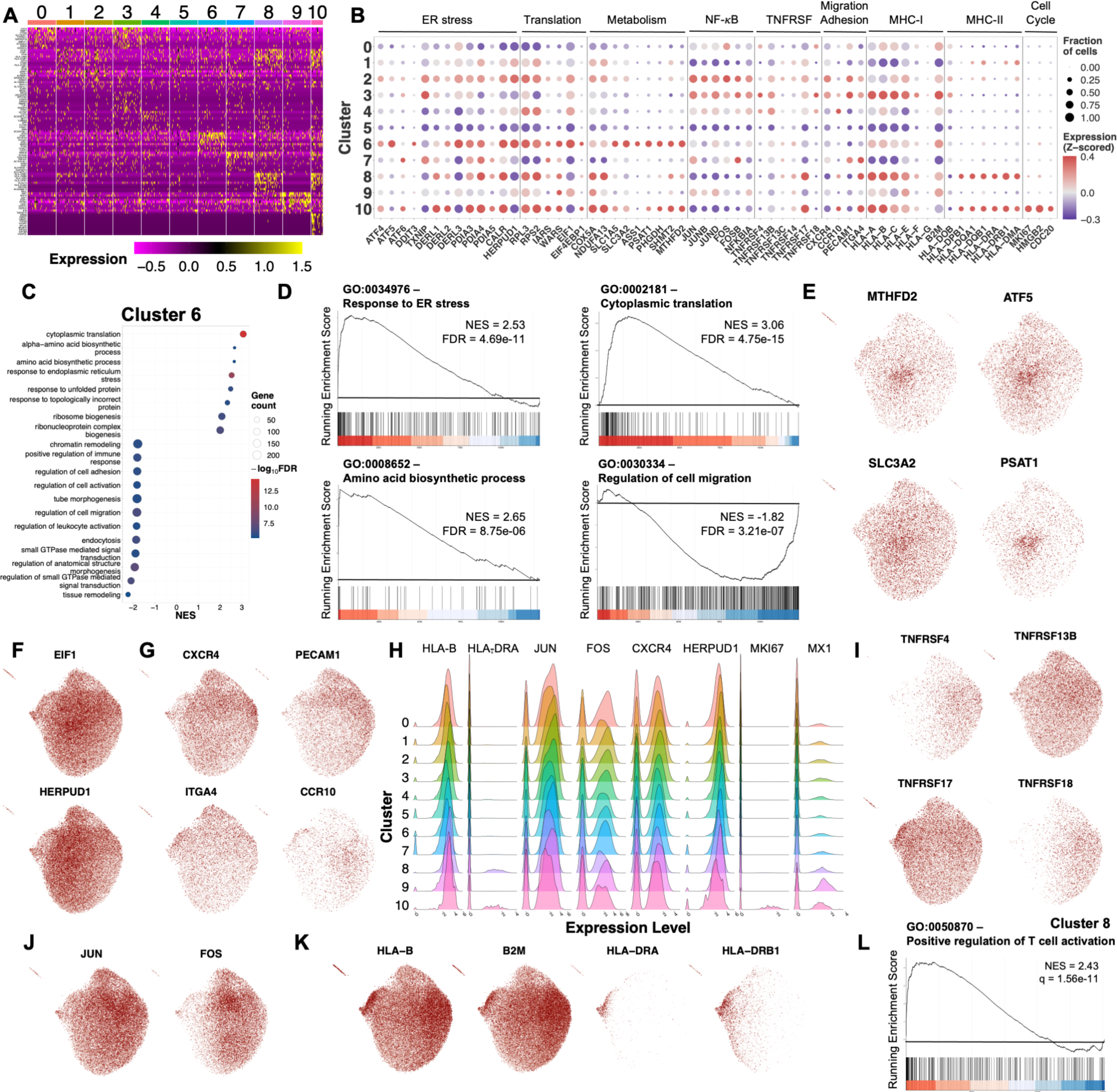
Transcriptional features of human bone marrow plasma cell subsets. (**A**) Heatmap of the top differentially expressed genes between clusters. (**B**) Bubble plot depicting expression of select genes associated with biological functions relevant to plasma cell biology. Bubble color represents expression compared to the mean. Bubble size represents the fraction of cells expressing the gene. (**C-D**) Gene set enrichment analysis (GSEA) of GO gene sets in cluster 6. In C, bubble color represents false discovery rate (FDR) and bubble size represents number of genes found in the dataset that belong to the gene set. NES: Normalized Enrichment Score, q = False Discovery Rate. (**E-K**) Expression of genes associated with amino acid and folate metabolism (E), ER stress (F), migration and adhesion (G), TNF superfamily receptors (I), NF-κB signaling (J), MHC antigen presentation (K). (**L**) Enrichment plot for GSEA performed on Cluster 8.

As previously reported (*5, 35*), we observed differences in the expression of TNF receptor superfamily genes and NF-κB signaling (Fig. 2H-J). However, their expression did not seem to be correlated. Unexpectedly, we identified MHC-I genes to be differentially expressed among mature BMPCs (Fig. 2K). Cluster 8 showed marked enrichment of MHC-II genes (Fig. 2K) and GSEA identified gene sets associated with the regulation of T cell activation as enriched within cluster 8 (Fig. 2L), which supports that these BMPCs have an immature phenotype (*36*).

GSEA was consistent with an interferon signature in cluster 9 and an active cell cycle signature for cluster 10. It also identified differential regulation of mitochondrial energy production (up in clusters 0, 1, 5, and 8; down in clusters 2, 3, and 7), ribosome synthesis (up in clusters 1 and 4; down in clusters 3 and 7), translation (up in cluster 5; down in cluster 3), ROS detoxification (up in cluster 0), T cell activation (up in cluster 8; down in clusters 0, 4, and 6), production of TNFSF cytokines (up in cluster 4), and retrograde transport of proteins from Golgi and ER (down in cluster 4) (Fig. S4).

Our results shed light on the heterogeneity of BMPCs beyond IgH expression. In addition, our dataset confirms that there is heterogeneous NF-κB signaling and TNFRSF expression among BMPCs, as has been found in other datasets (*5*). We also identify ER stress, MHC expression levels, and metabolism-associated genes as additional sources of BMPC heterogeneity.

### Origins of human BMPCs

BMPCs have been proposed to progressively lose CD19 expression as they mature into long-lived cells (*4, 5*). We observed heterogeneous surface CD19 expression among mature BMPCs (clusters 2 & 3). Mouse BMPC longevity has been shown to be correlated with different surface markers, such as TIGIT, CXCR3, EpCAM, and Ly6A (*32*). However, we did not observe differences in expression of these markers among human BMPCs. Therefore, we analyzed SHM levels and clonal dynamics with respect to CD19 protein levels.

We observed that SHM levels are higher in CD19^high^ BMPCs and lower in CD19^low^ BMPCs (Fig. 3A-B). This held true for IgA and IgG BMPCs, while IgE, IgD, and IgM BMPCs with different surface CD19 expression levels did not exhibit differences in IgH SHM (Fig. S5). In addition, BMPCs with higher SHM are more likely to be CD19^high^ than BMPCs with low SHM and vice versa (Fig. 3C), and we found a significant correlation between surface CD19 expression and IgH SHM (Fig. S5). IgG and IgM BMPCs were more prevalent in the CD19^low^ population, while IgA BMPCs were more prevalent in the CD19^high^ BMPC population (Fig. 3D), which was consistent across individuals (Figs. S6-7).

**Fig. 3.**
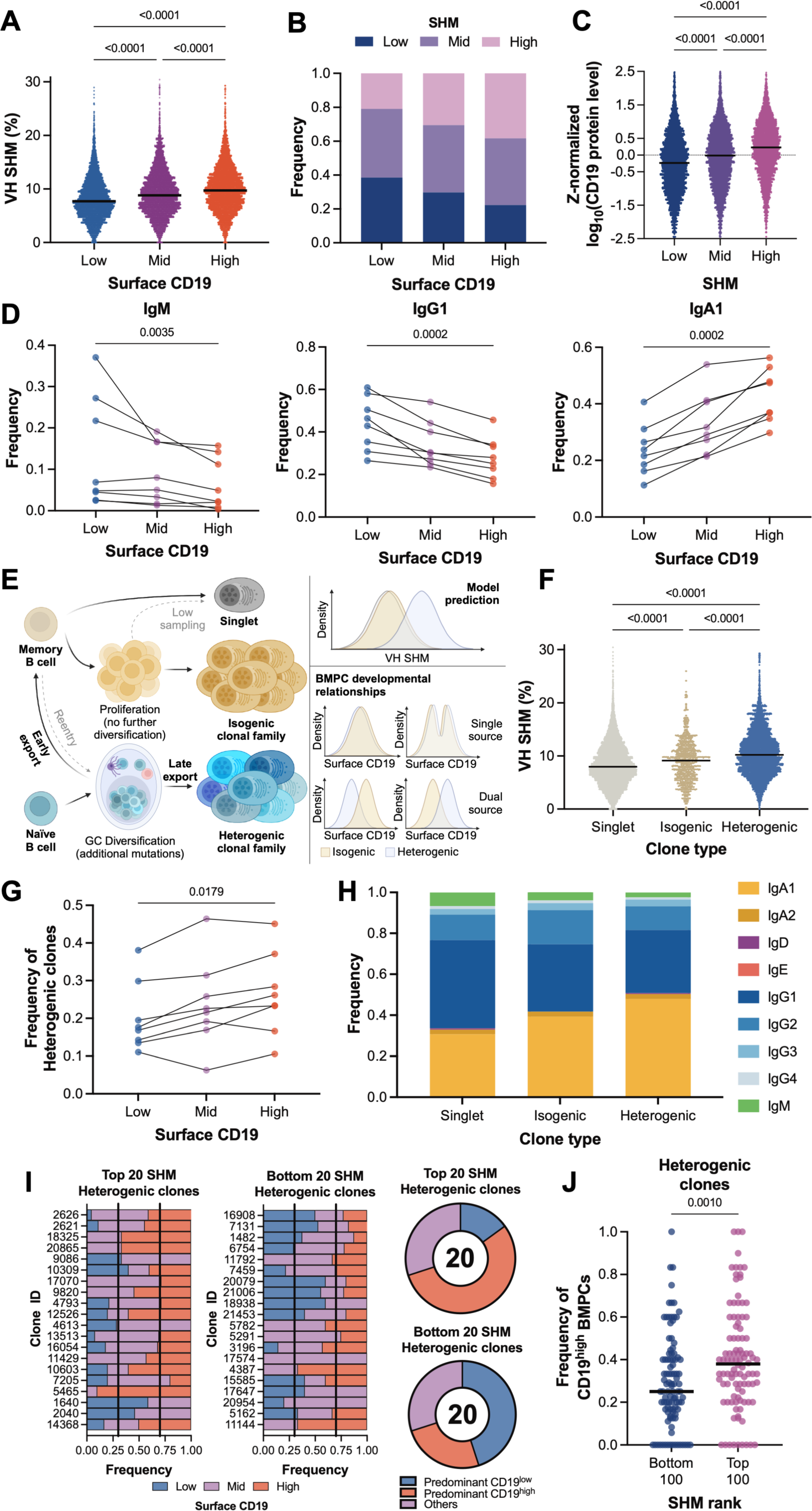
Surface CD19 expression suggests differential bone marrow plasma cell origins in humans. (**A**) IgH somatic hypermutation (SHM) among BMPCs categorized by surface CD19 protein expression (30% lowest, 40% medium, 30% highest). Kruskal-Wallis test was used to assess statistical differences. (**B**) Frequency of BMPCs with different IgH SHM levels (30% lowest, 40% medium, 30% highest) among surface CD19 protein expression categories. (**C**) Normalized surface CD19 protein expression levels among IgH SHM categories. Kruskal-Wallis test was used to assess statistical differences. (**D**) Frequency of IgM, IgG1, and IgA1 BMPCs among surface CD19 protein expression categories. Lines connecting dots represent an individual donor. Friedman test was used to assess statistical differences. (**E**) Schematic depicting the proposed relationship between clonal structure and BMPC source. (**F**) IgH SHM among different clonal structures. Kruskal-Wallis test was used to assess statistical differences. (**G**) Frequency of heterogenic clones among surface CD19 protein expression bins. Kruskal-Wallis test was used to assess statistical differences. (**H**) IgH isotypic composition among clone structures. (**I**) Surface CD19 expression bins among the 20 most mutated and 20 least mutated heterogenic clonal families. Vertical black lines represent expected proportions based on CD19 bin definition. Clonal families were defined as predominantly CD19^low^ if more than 30% of the family is CD19^low^ and less than 30% of the family is CD19^high^, and vice versa. (**J**) Frequency of CD19^high^ BMPCs among the 100 most mutated and 100 least mutated heterogenic clonal families. Mann-Whitney U test was used to assess statistical differences.

If CD19^low^ BPMCs mature from CD19^high^ BMPCs, it is unclear why they would have lower SHM. One possibility is that BMPCs with higher SHM levels may have higher attrition rates due to potentially higher genotoxic stress induced by off-target mutations, as has been suggested for memory B cells (*37*). A non-mutually exclusive possibility is that CD19^low^ and CD19^high^ cells descend from different cellular ancestors. Essentially all BMPCs have undergone substantial SHM (Fig. S2F), suggesting passage through the GC B cell stage during their developmental path. GC B cells can develop directly into plasma cells after primary antigen exposure or differentiate into circulating memory B cells (MBCs), which can subsequently differentiate into plasma cells after a recall stimulus. We surmised that aspects of intra-clonal SHM could help distinguish from which of these scenarios BMPCs arose (Fig. 3E). A key assumption in this model is that MBCs and GC B cells (or their descendants) have different rates of CD19 loss upon differentiation.

Long-lived GC-derived MBCs possess lower levels of SHM than GC-derived plasma cells do because MBCs exit from GCs earlier than plasma cells do (*38*). Circulating MBCs rapidly proliferate and differentiate into antibody secreting cells upon secondary antigen exposure with minimal re-entry to GCs (*39*). Therefore, we expect MBC progeny to be either singlets or identical clones of each other (isogenic clones), reflecting MBC-to-plasma cell differentiation in the absence of GC re-entry. On the other hand, we expect direct GC B cell-to-plasma cell exports to have a relatively higher degree of intra-clonal variations of the same sequence (heterogenic clones), given GC dynamics (Fig. 3E). Indeed, heterogenic clones show higher SHM levels than isogenic clones do (Fig. 3F), supporting the idea that MBC recall may contribute to the CD19^low^ BMPC pool, and direct GC B cell to plasma cell differentiation may contribute to the CD19^high^ BMPC pool. Consistent with this hypothesis, heterogenic clones tended to accumulate in the CD19^high^ BMPC compartment (Fig. 3G). Furthermore, the frequency of CD19^low^ BMPC-associated IgG and IgM clones was also higher among singlets and isogenic clones, consistent with abundances of long-lived peripheral IgM and IgG MBCs (*40*). The frequency of CD19^high^ BMPC-associated IgA was higher among heterogenic clones (Fig. 3H), consistent with high levels of IgA class switching and direct plasma cell outputs from mucosal tissue GCs (*40, 41*).

To further dissect the relationship between surface CD19 expression and BMPC origin, we looked at the composition of the most mutated and least mutated heterogenic clones (Fig. 3I & S8). As expected, CD19^high^ BMPCs tended to be overrepresented among the most mutated heterogenic clones, while CD19^low^ BMPCs tended to be overrepresented among the least mutated heterogenic clones (Fig. 3I-J). All types of clonal structures were homogenously distributed among clusters (Fig. S8).

Taken together, our data are consistent with a model where direct GC-derived plasma cells contribute to the CD19^high^ BMPC pool, while plasma cells derived from MBCs that directly differentiate into plasma cells contribute to the CD19^low^ BMPC pool. The differences observed for surface CD19 expression among BMPCs could be explained by different rates of CD19 expression loss and/or rates of removal of CD19 protein from the cell surface from MBC or GC B cells. This model provides a mechanistic framework that is consistent with observations that recalled immunity induces longer-lived, stable antibody levels (*6, 7*), and is non-mutually exclusive with a model in which CD19^high^ BMPCs lose CD19 expression as they mature into CD19^low^ BMPCs (*5*).

### IgE BMPCs in food-allergic individuals

It has long been thought that long-lived plasma cells in the bone marrow may contribute to allergic sensitization, particularly in peanut allergy, where sensitization can last for decades without new allergen exposure (*21, 22*) Instances of transfer of food allergies through bone marrow donations have also been documented, which would be consistent with a long-lived source of allergen-specific antibodies (*42, 43*). However, allergen-specific BMPCs are not readily detected due to their extreme rarity.

By combining VDJ and 5’ transcriptome reads, we identified 54 IgE BMPCs in our dataset (Fig. 4A & S9). Even though 49% of BMPCs were derived from food-allergic individuals, 47/54 (87%) of IgE BMPCs were derived from these individuals, who also had significantly higher IgE titers in blood (Fig. S10). We also found a larger than expected fraction of IgM and IgD BMPCs, and a substantially lower fraction of IgG4 BMPCs in food-allergic individuals (Fig. 4B). These proportions also held true when donors were analyzed separately (Fig. S10). The inverse correlation of IgG4 BMPCs in peanut allergy is consistent with the notion that IgG4 may offer a protective shield to reduce clinical sensitization (*44–46*). Neither sex nor age were correlated with fraction of BMPCs of any given isotype, except maybe for IgM, for which age tended to be correlated with a lower proportion of IgM BMPCs (P = 0.0526) (Fig. S10).

**Fig. 4.**
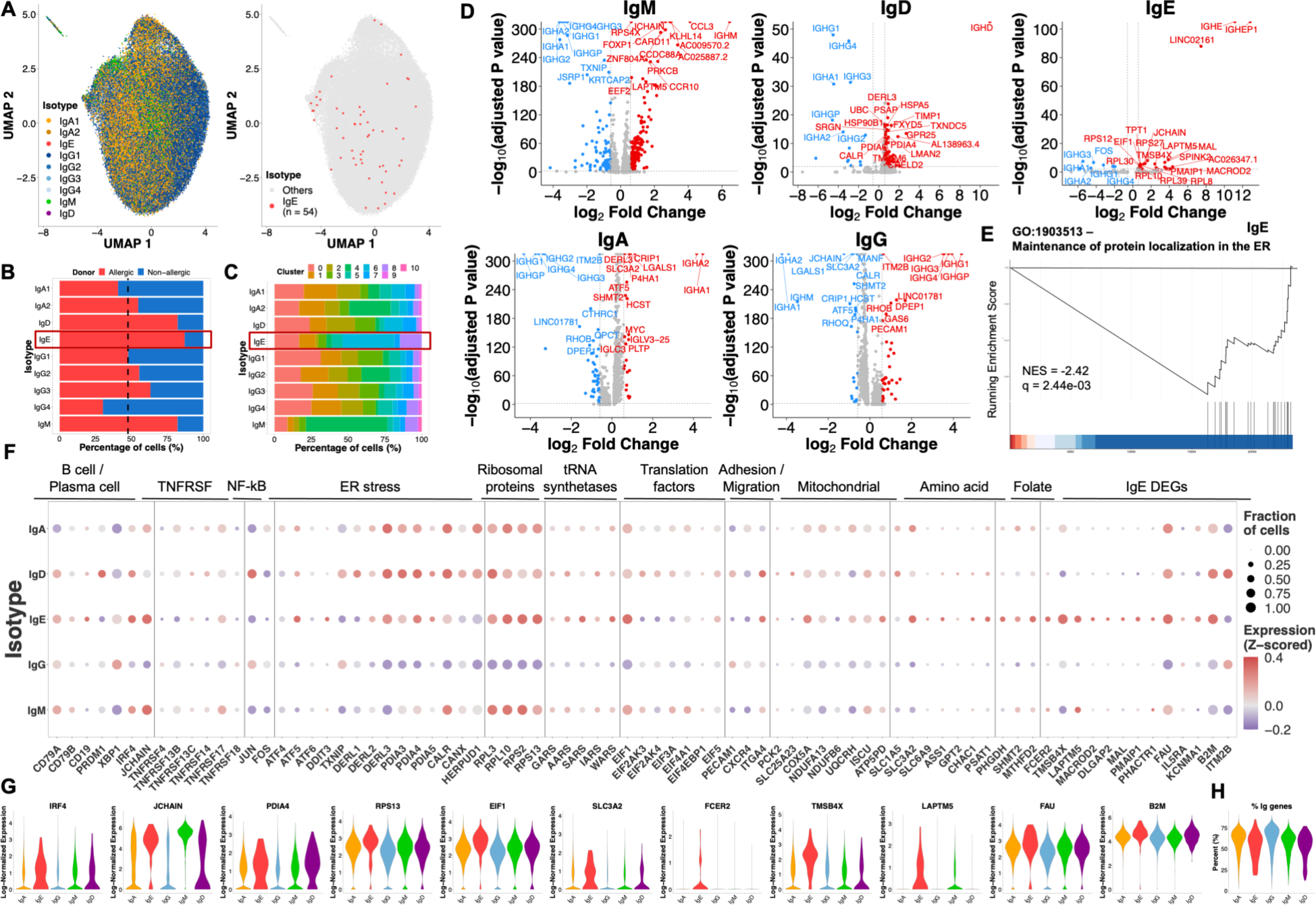
IgE bone marrow plasma cells show signs of high metabolic demand and ER stress. (**A**) UMAP projection of cells, colored by IgH isotype (left) and highlighting IgE BMPCs (right). (**B-C**) Contributions of food allergic or non-allergic donors (B) and cluster identities (C) to BMPCs of different IgH isotypes. (**D**) Volcano plots for comparisons between BMPCs of a given isotype versus all other BMPCs. Isotypes with multiple subclasses were aggregated into a single isotype class. (**E**) Enrichment plot for IgE cells. (**F-G**) Gene expression separated by isotype class. Bubble color represents expression compared to the mean. Bubble size represents the fraction of cells that express the gene. (**H**) Percentage of immunoglobulin genes expressed by BMPCs of different isotypes.

The enrichment of IgE BMPCs in cluster 6, and to a lesser extent in cluster 8 (Fig. 4C & S10), pointed towards an active metabolism/ER stress and immature phenotype, respectively. Most IgE BMPC sequences consisted of singlets, but we found an IgE clone that was composed of 2 non-identical sequences, as well as a clone composed of one IgG1 and one IgE non-identical sequences (Fig. S11). These data are consistent with both the rarity of IgE BMPCs and an MBC source of IgE BMPCs based on low surface CD19 expression (Fig. S8B) and a bias towards intermediate-low SHM levels (90.1% of IgE BMPCs with VDJ data).

Analysis of IgH isotype-specific DEGs (Fig. 4D) showed that IgM cells expressed the highest amount of *JCHAIN*, which is consistent with IgM’s pentameric structure. They also upregulated chemokine receptors associated with skin and gut homing (e.g. *CCR10*) (*35*), downregulated genes associated with ER stress (e.g. *TXNIP*), and upregulated genes associated with a more immature phenotype (*FOXP1, KLHL14*). IgA BMPCs expressed high levels of genes associated with amino acid metabolism (*SLC3A2, ATF5, SHMT2*), as well as higher levels of *MYC*. This finding is consistent with a model in which IgA BMPCs are exported from the gut as GC emigrants rather than from a memory B cell pool. IgE BMPCs expressed high levels of *LAPTM5*, a known negative regulator of BCR signaling. GSEA revealed that genes associated with the maintenance of proteins in the ER lumen are downregulated in IgE BMPCs (Fig. 4E & S11). Both IgD and IgE BMPCs expressed high levels of ER stress- and translation-associated genes (*PDIA4, PDIA5, CALR, DERL3*, ribosomal proteins, tRNA synthetases, translation initiation factors) (Fig. 4F-G). IgE BMPCs also upregulated genes associated with plasma cell identity transcription factor *IRF4*, the amino acid transporter *SLC3A2*, and actin sequester protein *TMSB4X*. While allergen-specific MBCs that differentiate towards IgE-producing cells have been recently described to express CD23, IL4R, and IL13R (*12, 13*), we only found slight differences in *FCER2* expression (coding for CD23) on IgE BMPCs. Finally, IgE BMPCs expressed very high levels of *JCHAIN*, which were higher than those of IgA BMPCs and second only to those of IgM BMPCs (Fig. 4F-G & S11).

Given that *JCHAIN* expression has been linked to high Ig production independent of IgH isotype (*47–50*), we decided to explore patterns of *JCHAIN* expression in our dataset. We found that CD19^low^ BMPCs and SHM^low^ BMPCs expressed higher amounts of *JCHAIN*, independently of IgH isotype (Fig. S12). Additionally, heterogenic clonal families expressed slightly lower amounts of *JCHAIN* compared to singlets and isogenic clonal families (Fig. S12). These results point towards a high Ig secretion phenotype for MBC-derived BMPCs. On the other hand, it might also be reflective of a mucosal origin, since *JCHAIN* expression in IgG and IgD mucosal plasma cells has been noted (*48, 51–53*). We do not exclude the possibility of an entirely unknown function, given that *JCHAIN* expression has been observed in other immune and non-immune cell types in both vertebrates and invertebrates (*54–57*).

Our results are consistent with a high Ig production phenotype and/or a mucosal origin for IgE BMPCs. However, immunoglobulin variable region transcripts did not seem to vary appreciably among BMPCs of different isotypes (Fig. 4H), so additional post-transcriptional mechanisms could potentially contribute to the phenotype.

### BMPCs can be a source of peanut-binding IgE antibodies

To test for IgE BMPCs’ antigen specificity and antibody function, we cloned IgE BMPC sequences from our initial cohort, one additional peanut-allergic individual, and an individual with a dairy food allergy. These two subjects did not pass our criteria to be included in gene expression analyses due to strong batch effects, but BCR sequence data were available, nonetheless. All VDJ sequences were cloned into a human IgG1 backbone to screen for peanut reactivity. Out of 32 cloned IgE BMPC sequences derived from food-allergic individuals, we found two peanut-reactive, IgE BMPC-derived monoclonal antibodies from two donors with history of anaphylaxis against peanuts (P0069 and P0544) (Fig. 5A). While antibody P69P4 strongly bound to Ara h 2 but not to Ara h 6 peanut antigens, antibody P544P1 did not bind to Ara h 2 and weakly bound to Ara h 6, as determined by ELISA (Fig. 5A). Neither antibody sequence was polyreactive, as determined by ELISA binding to Spike protein or mouse serum albumin (MSA), nor were they previously reported public peanut-binding sequences (*10, 17*).

**Fig. 5.**
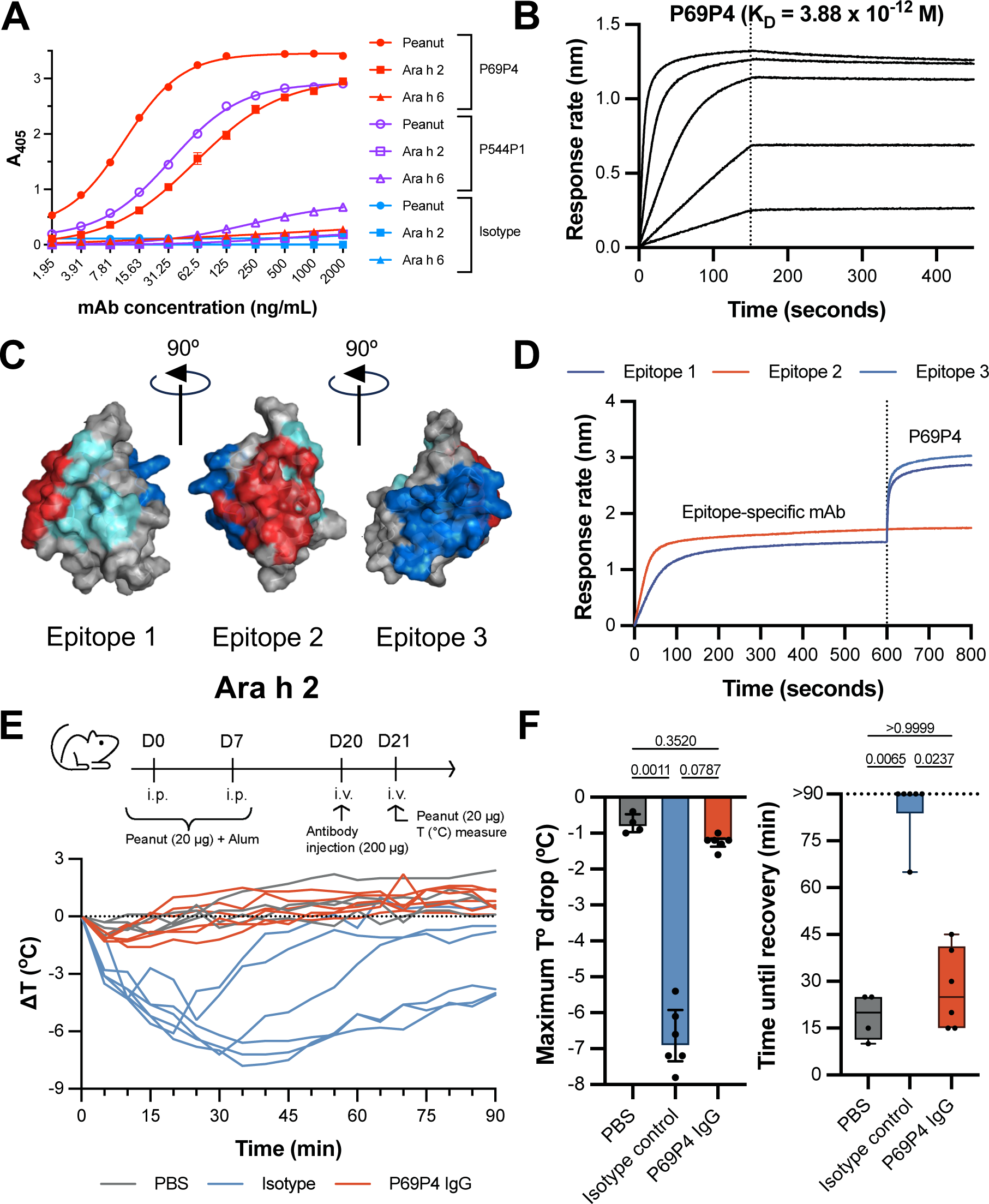
IgE human bone marrow plasma cells harbor allergen-specific antibody sequences. (**A**) ELISA testing for binding of P69P4 in a human IgG1 backbone (red) to peanut extract, Ara h 2, and Ara h 6. An anti-SARS-CoV-2 Spike IgG antibody was used as an isotype control (blue). (**B**) Affinity curves to biotinylated natural Ara h 2 by BLI for P69P4. (**C**) Molecular modeling of Ara h 2 (PDB: 8G4P). Epitopes 1, 2, and 3 determined in LaHood *et al.* (*58*) are highlighted in teal, red, and dark blue, respectively. (**D**) Epitope binning sensograms for P69P4 using primary antibodies, 23P34 (Epitope 1.1), 13T5 (Epitope 2), and 22S1 (Epitope 3) from LaHood *et al.* (*58*). Inhibition of response increase indicates overlapping epitopic regions targeted. (**E-F**) Temperature curves and quantification of anaphylactic responses of peanut-sensitized BALB/c mice after antibody injection and intravenous peanut challenge. PBS mice were not sensitized to peanut or administer mAbs prior to i.v. peanut challenge. An anti-SARS-CoV-2 Spike IgG antibody was used as an isotype control. Kruskal-Wallis test was used to assess statistical differences.

We further characterized the peanut-binding monoclonal antibodies through biolayer interferometry (BLI). While P544P1 did not bind specifically to biotinylated Ara h 2 or Ara h 6 in this assay, P69P4 showed a sub-nanomolar affinity (K_D_) for biotinylated natural Ara h 2 (Fig. 5B). Competition-based epitope mapping of this monoclonal antibody identified binding localized primarily to helix 3 of Ara h 2 (Epitope 2, PDB 8G4P) (*58, 59*) (Fig. 5C-D). Notably, subject P0069, from whom the antibody was derived, underwent oral immunotherapy for peanut allergy in the past. Antibodies that bind epitope 2 in Ara h 2 have been associated with positive responses to oral immunotherapy. However, she failed to induce tolerance against peanut antigens during therapy.

To test whether this IgE BMPC-derived antibody could have physiologically relevant binding to peanut *in vivo*, we sensitized BALB/c mice with peanut extract. We then administered P69P4 monoclonal antibody (P69P4 IgG) or an isotype-matched mAb (isotype) the day before challenging with intravenous peanut extract to induce anaphylaxis (Fig. 5E). Given that P69P4’s VDJ sequence (IgE BMPC-derived) is expressed on an IgG1 backbone, we expected that it would bind to peanut antigens but would not activate FcεRI receptors, therefore effectively blocking any anaphylactic responses upon peanut challenge. As expected, mice administered with the IgG isotype control developed anaphylactic responses, as measured by a drop in body temperature after peanut challenge. However, mice that were administered P69P4 prior to challenge did not exhibit a drop in body temperature, responding indistinguishably from unsensitized mice (Fig. 5E-F). While the presence of only two instances of peanut-binding IgE plasma cells, each identified in independent peanut allergic study subjects within this cohort, may be limited, it does indicate the potential for IgE BMPCs in peanut-allergic individuals to generate peanut-binding antibodies. Notably, one of these instances demonstrates significant physiological relevance *in vivo*.

In summary, we show that human BMPCs exhibit intrinsic transcriptional heterogeneity with different cellular origins. Recalled MBCs may contribute directly to longer-lived CD19^low^ BMPCs while GC-derived plasma cells may contribute relatively more to the CD19^high^ BMPC pool. Together with the idea that CD19^low^ BMPCs contain the longest-lived cells, this model is compatible with observations showing that immune recall induces greater stable antibody levels and with the notion that loss of CD19 expression during BMPC maturation occurs independently of cellular ancestry. This work also highlights J chain expression as a potential indicator of greater per cell antibody production, enriched in longer lived CD19^low^ BMPCs, along with relatively lower SHM on average. In addition, this work also shows a high metabolism/ER-stress cellular state in IgE BMPCs and identifies them as a source of allergen-specific IgE in peanut allergy, with functional consequence *in vivo*. The data here shed light on sources of pathogenic antibodies in food allergy and cellular ancestry of human bone marrow plasma cells.

## Acknowledgments

We acknowledge Moshe Ben-David and Yanay Ofran for Ara h 2 and Ara h 6 proteins. Michel Nussenzweig for expression plasmids. We thank Stephanie Eisenbarth, Emily Sincalco, Hans Oettgen, Renan de Carvalho, Courtney Stump, and Sarah Kate Lane-Reticker for their helpful comments during manuscript preparation.

## Funding

National Institutes of Health grant R01AI158811 (DRW)

National Institutes of Health grant R01AI170715 (DRW)

National Institutes of Health grant P01AI156072 (DRW)

Food Allergy Science Initiative (FASI)

Sanofi (DRW)

Anonymous donor (DRW)

## Author contributions

Conceptualization: GAP, VR, PT, DRW

Data curation: GAP, VR, SS, ASM, RE, AA, AP, DRW

Formal analysis: GAP, VR, DKY, SS

Funding acquisition: SUP, DRW

Investigation: GAP, VR, DKY, SS, SK, PT, OMR, ZA, DRW

Methodology: GAP, VR, SS, OMR

Project administration: GAP, DRW

Resources: OMR, SUP

Supervision: GAP, DRW

Visualization: GAP, VR, DKY, SS, OMR

Writing – original draft: GAP, DRW

Writing – review & editing: GAP, VR, DKY, SS, PT, ZA, DRW

## Materials and Methods

### Materials

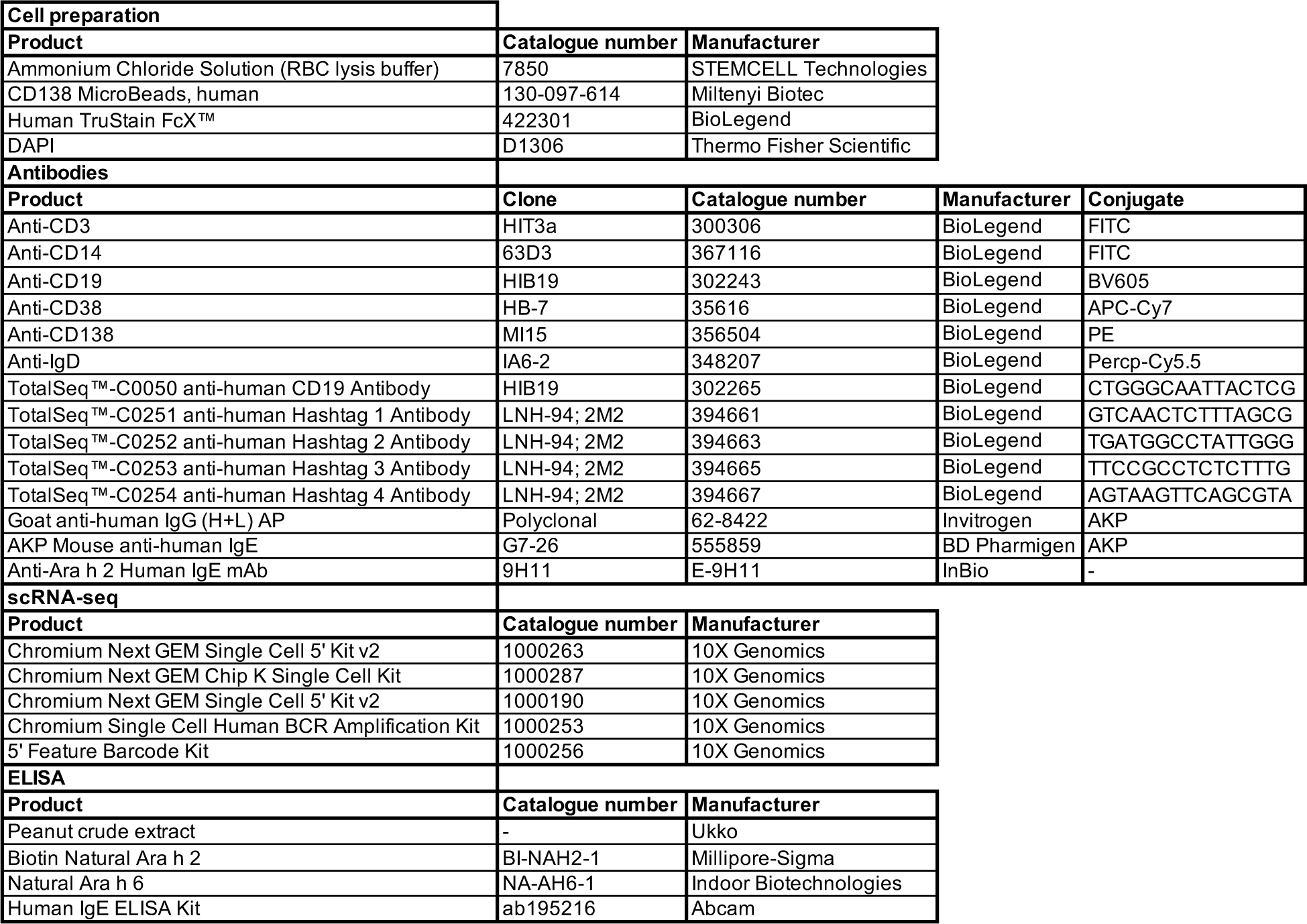

### Bone marrow aspirates, peripheral blood collection, and sample preparations

All clinical procedures were approved by the Institutional Review Board of Brigham and Women’s Hospital (IRB Protocol Number: 2018P002007). Participants provided written consent for the study, we conducted laboratory tests to gather clinical data, and stored the data in a secure data repository. We recruited study participants who either had a history of food allergy and anaphylaxis or not (Supplementary Table 1). Bone marrow (BM) aspirates and peripheral blood were collected from each group. We obtained 20-40 mL of BM aspirate from the posterior superior iliac spine using heparin or EDTA collection tubes (BD). Red blood cells (RBCs) were lysed in BM samples using 0.8% ammonium chloride solution (Stem Cell Technologies). Samples were then directly used for downstream applications, including magnetic enrichment, flow cytometry, cell sorting, and single B cell sequencing. Peripheral blood was collected on EDTA tubes (BD Biosciences). Peripheral blood mononuclear cells (PBMCs) were isolated from peripheral blood by density gradient centrifugation using Ficoll PLUS medium (Cytiva) and stored in serum-free cryopreservation medium Cellbanker 2 (Amsbio). Plasma was saved after Ficoll gradient separation and stored at −80°C until further use. PBMCs were aliquoted and stored until use at −80°C. RBCs were lysed using RBC lysis buffer (eBioscience) before downstream applications after thawing.

### Bone marrow plasma cell sorting

RBC-lysed bone marrow cells were positively selected using CD138 microbeads kit (Miltenyi Biotec) and stained with anti-CD138-PE (clone MI15, BioLegend, 1:50), anti-CD38-APC-Cy7 (clone HB-7, BioLegend, 1:50), anti-CD19-BV605 (clone HIB19, BioLegend, 1:50), anti-IgD-PerCP-Cy5.5 (clone IA6-2, BioLegend, 1:50), anti-CD3-FITC (clone HIT3a, BioLegend, 1:50), anti-CD14-FITC (clone 63D3, BioLegend, 1:50), and DAPI (ThermoFisher Scientific, 1:2,000) (see Methods). Additionally, sorted samples were labeled with TotalSeq-C Hashtag 1–5 antibodies (TotalSeq-C0251, TotalSeq-C0252, TotalSeq-C0253, TotalSeq-C0254, TotalSeq-C0255) (0.5 µg per 1,000,000 cells) and TotalSeq-C0050 anti-CD19 antibody (BioLegend) for demultiplexing individual samples in downstream analysis. CD38^hi^CD138^+^ cells were sorted with a FACSAria II cell sorter (BD Biosciences) into cooled 1.5-mL tubes (BioRad). When TotalSeq-C0050 anti-CD19 antibody was not available, cells were further sorted into CD19^high^ or CD19^low^ populations.

FACS data were collected with the BD FACSDiva (v.8.0) software. FlowJo v.10.7.1 (BD Biosciences) was used for flow cytometry data analysis.

### 10X Genomics scRNA-seq

Using a Single-Cell 5′ Library and Gel Bead kit v.1.1 (10X Genomics, 1000165) and Next GEM Chip G Single-Cell kit (10X Genomics, 1000120), the magnetically enriched, FACS-sorted BMPC single cell suspension was loaded onto a Chromium single-cell controller (10X Genomics) to generate single-cell gel beads in the emulsion (GEMs) according to the manufacturer’s protocol. 4,000 – 15,000 cells were added to each channel and approximately 50 – 70 % were recovered (Supplementary Table 2). Captured cells were lysed and released RNA was barcoded through reverse transcription in individual GEMs. The 5′ gene expression (GEX) libraries, single-cell V(D)J libraries (1000016) and cell surface protein libraries were constructed according to manufacturer protocols. Library quality was assessed using a 2200 TapeStation (Agilent). mRNA libraries were sequenced to an average of 30,000 reads per cell using Illumina Novaseq sequencer with a paired-end 150-bp (PE150) reading strategy (Dana Farber Cancer Institute).

### 10X data pre-processing and quality control

scRNA-seq reads were processed with Cell Ranger (v3.1), which quantified transcript counts per putative cell. Quantification was performed using the STAR aligner against the GRCh38 transcriptome.

Data was analyzed with the Seurat package (v5.0.1) (*60*) in RStudio. Cells expressing less than 5% of mitochondrial genes and between 200 and 4,000 RNAs were retained for further analysis. Barcodes were demultiplexed based on hashtag oligonucleotide count distribution and proportion among all hashing antibodies in a run-dependent manner. Cells expressing *CD3E* (T cells), *NKG7*, *PRF1* (NK cells), *S100A9* (neutrophils), *FCGR3A*, *CD14* (monocytes), *COL1A1*, *APOE*, *CSFR3* (macrophages and fibroblasts), *HBB* (erythroid) were discarded.

### Isotype assignation of cells

Isotype assignation was performed via a combination of gene expression (GEX) and BCR (VDJ) libraries. If a barcode in GEX had a corresponding barcode in VDJ with heavy chain isotype information, that barcode was assigned the identified VDJ isotype. If a barcode in GEX had a corresponding barcode in VDJ without heavy chain isotype information, or if it did not have a corresponding barcode in VDJ, heavy chain isotype was assigned based on GEX via the following criteria: i) more than half of IgH isotypes corresponded to a given isotype, and ii) the cell expressed more than 20 counts of the given isotype. Cells that were not assigned a heavy chain isotype with confidence were discarded. When benchmarked against VDJ-based isotype assignation, GEX-based isotype inference performed accurately for 95.09-99.99% of the barcodes with corresponding VDJ isotype information (Fig. S9). This method identified isotype class and subclass with remarkable specificity, ranging from 95.56% to 99.99% depending on heavy chain isotype (Fig. S9). Isotype class (IgM/D/E/G/A & IgK/L) was defined based on isotype subclass determined by GEX or VDJ data (IgM/D/E/G1/G2/G3/G4/A1/A2 & IgK/L1/L2/L3/L4/L5/L6/L7).

### Batch-correction, clustering and surface CD19 protein expression

Variable feature selection was performed after excluding immunoglobulin genes (V, D, J, and C genes for all IgH and IgL isotypes, including pseudogenes, and contaminant non-annotated Ig genes *AC233755.1* and *AC233755.2*) and *XIST* to account for sex differences. PCA was performed on the initial dataset containing Ig genes based on the variable features defined without Ig genes.

Batch correction between 10X runs was performed with Harmony (v1.2.0) (*61*). Cells were clustered using the Louvain algorithm with a resolution of 0.67. Integration metrics were calculated using the scIntegrationMetrics package (v1.1) (*62*). Runs that did not at least double their iLISI score after batch correction were not included in clustering and gene expression-related analyses (see Supplementary Table 1) but were retained for IgE BCR sequence analysis (see “Antibody production and binding reactivity screening with ELISA” section below).

Surface CD19 expression was assessed using oligonucleotide-tagged anti-CD19 antibodies (see “Bone marrow plasma cell sorting” and “Materials” sections above). Antibody capture data was log-transformed and Z-scored to normalize for batch effects between runs. CD19 expression bins (low, mid, high) were defined by ranking surface CD19 expression: the top 30 expressors were defined as CD19^high^, the middle 40 were defined as CD19^mid^, and the bottom 30 were defined as CD19^low^.

### Differential gene expression testing and gene set enrichment analysis (GSEA)

Differentially expressed genes (DEGs) were identified on whole data using FindMarkers in Seurat. DEGs were identified for clusters and for isotype classes by comparing each group versus all the rest. For IgE cells, additional DEGs were identified by comparing IgE cells versus other heavy chain isotypes on a 1 vs 1 basis. Data from these analyses were used for heatmaps, bubble plots, violin plots, and volcano plots.

Prior to GSEA (*63*), data were pseudobulked with Seurat (v5.0.1) by donor and by cluster, or by donor and isotype class. DEGs were determined using DESeq2 (v1.42.0) (*64*) and these data were used as input for GSEA using clusterProfiler (v4.10.0) (*65*).

### Somatic hypermutation and clonal composition analyses

Input VDJ data was processed using IgBlast to determine gene segment usage and CDR positions. Somatic hypermutation (SHM) was determined by comparison of the BCR heavy chain nucleotide sequence with the germline sequence. SHM bins (low, mid, high) were defined by ranking heavy chain SHM: the top 30 most mutated cells were defined as SHM^high^, the middle 40 were defined as SHM^mid^, and the bottom 30 were defined as SHM^low^.

The Immcantation package was then used to define BCR clonotypes (Change-O v1.3.0) (*66*) based on heavy chain CDR3 amino acid sequence similarity and identical usage of IGHV and IGHJ genes. The hamming distance cut-off (0.85) was determined by SHazaM (v1.1.2) (*66*). Clonal relationships were graphed using circlize (v0.4.15) (*67*) in R.

Clonal families were classified into 3 categories based on their lineage structure – i) heterogenic clones, consisting of clones with at least 2 cells with unique sequences representing an expanded clone differing in mutations, ii) isogenic clones, consisting of clonotypes with at least 2 cells, where all cells have identical sequences, representing an expanded clone with identical mutations and iii) singlets, consisting of cells that are not part of an expanded clone and have unique sequences. Heterogenic and isogenic clonal families with at least 5 cells were used for CD19 distribution, SHM distribution, and isotype frequency analyses.

### Antibody production and binding reactivity screening with ELISA

Heavy and light chain variable domain gene fragments were synthesized from IgE BMPC scVDJ-seq data based on raw matrices. Antibody sequences were then cloned into human IgG1 vectors, expressed, and purified by Twist Bioscience.

To evaluate the binding reactivity of antibodies from BMPCs, ELISA was carried out. Briefly, 96-well ELISA plates (ThermoFisher) were coated with 100 ng/well of peanut extract protein (Ukko), 100 ng/well Ara h 2 (Millipore Sigma), or 50 ng/well Ara h 6 (Ukko) in 50 µL coating buffer (0.1M NaHCO_3_/Na_2_CO_3_, pH = 9.5) at 4°C overnight. Plates were blocked with 150 μL of 5% BSA in PBS for at least 2 hours. 50 µL of purified antibodies were added to the plates (4 µg/mL – 0.0024 µg/mL, 2-fold serial dilutions in 2% BSA) and incubated for 1 hour. Plates were washed 4 times with PBS + 0.05% Tween-20 (PBST). 50 µL/well of anti-human IgG-alkaline phosphatase (Southern Biotech) (1 μg/mL in PBST + 1% BSA) was added and incubated for 1 hour at room temperature. Plates were washed three times with PBST. 100 μL/well of developing solution (0.1 M glycine, pH 10.4, 1 mM MgCl_2_, 1 mM ZnCl_2_, 1.6 mg/mL alkaline phosphatase substrate *p*-nitrophenyl phosphate (Sigma-Aldrich)) was then added to the plates and incubated for 2 hours at room temperature. Absorbance was measured at 405 nm using a microplate reader (Biotek Synergy H1). EC_50_ and AUC were calculated with GraphPad Prism 10.

### Quantification of mAbs and affinity determination

Purified IgG1 mAbs were quantified using anti–human Fab-CH1 (FAB2G) biosensors (Sartorius) as previously described (*58*). To measure affinity, biotinylated natural Ara h 2 (InBio) diluted in kinetics buffer (Dulbecco’s PBS, 1% BSA, 0.02% Tween-20) was loaded onto streptavidin sensors (Sartorius) for 100 seconds. Antibody affinities were measured using a minimum of 5 concentrations after 3-fold dilutions from 2 ug/mL. Experiments were performed using Octet R2 Protein Analysis System (Sartorius) and affinity was calculated using Octet Analysis Studio Software, version 12.2, and GraphPad Prism, version 9.3.1, as previously described.

### Epitope determination

To localize epitopes of mAbs, we used an in-tandem assay on BLI, as previously described (*58*). Briefly, antibodies were diluted in kinetics buffer, (10 μg/mL primary mAbs and 5 μg/mL secondary mAbs). For primary antibodies, we used previously described monoclonal Ara h 2 specific antibodies, 23P34, 13T5, 22S1, and 13T1, with known epitope characterization (PDBs 8DB4 and 8G4P) (*58, 59*). Streptavidin sensors (Sartorius) were loaded with biotinylated natural Ara h 2 (InBio). Secondary antibodies were grouped into a separate epitope if binding was 0.16 nm or higher (≥4 times the SD of nonbinding antibodies, based on previous work (*58*). Analysis was performed using Octet Analysis Studio Software, version 12.2, and GraphPad Prism, version 9.3.1.

### Protection of BALB/c anaphylaxis using allergen-specific IgG antibody

To evaluate *in vivo* binding of peanut-specific IgE BMPC BCR sequences (Fig. 5C), we employed an active peanut anaphylaxis model using female WT BALB/c mice (Jackson Laboratory, 000651). Mice were sensitized to peanut antigens by intraperitoneal (i.p.) injection at day 0 and day 7 with 20 µg peanut extract in 100 µL PBS mixed with 100 µL Alum (ThermoFisher). In parallel, mice immunized with PBS and Alum without peanut extract were prepared as a control group. On day 20, mice were intravenously (i.v.) injected with 200 µg of antibodies. On day 21, anaphylaxis was induced by challenging mice with i.v. injection of 20 µg peanut extract in 100 µL PBS. The mice’s temperature was measured with a DAS-8027IUS temperature probe system (BMDS) before challenge and every 5 minutes for 90 minutes after the antigen challenge.

### Statistical analysis and data visualization

Plots were generated in Prism Graphpad (v10.1.1) or in R using ggplot2 (v3.4.4) (*68*), circlize (v0.4.15) (*67*), Seurat (v5.0.1) (*60*), and clusterProfiler (v4.10.0) (*65*). Flow cytometry data was visualized using FlowJo v10.7.1.

Unpaired non-parametric tests were used to assess statistical differences. Mann-Whitney tests were used when comparing two independent groups, Kruskal-Wallis tests were used when comparing three or more independent groups, and Friedman tests were used when comparing three dependent groups. All *post-hoc* tests were performed by comparing all means or ranks against all other means or ranks. Sample sizes are indicated in figure legends.

## Supplementary Figures

**Figure S1.**
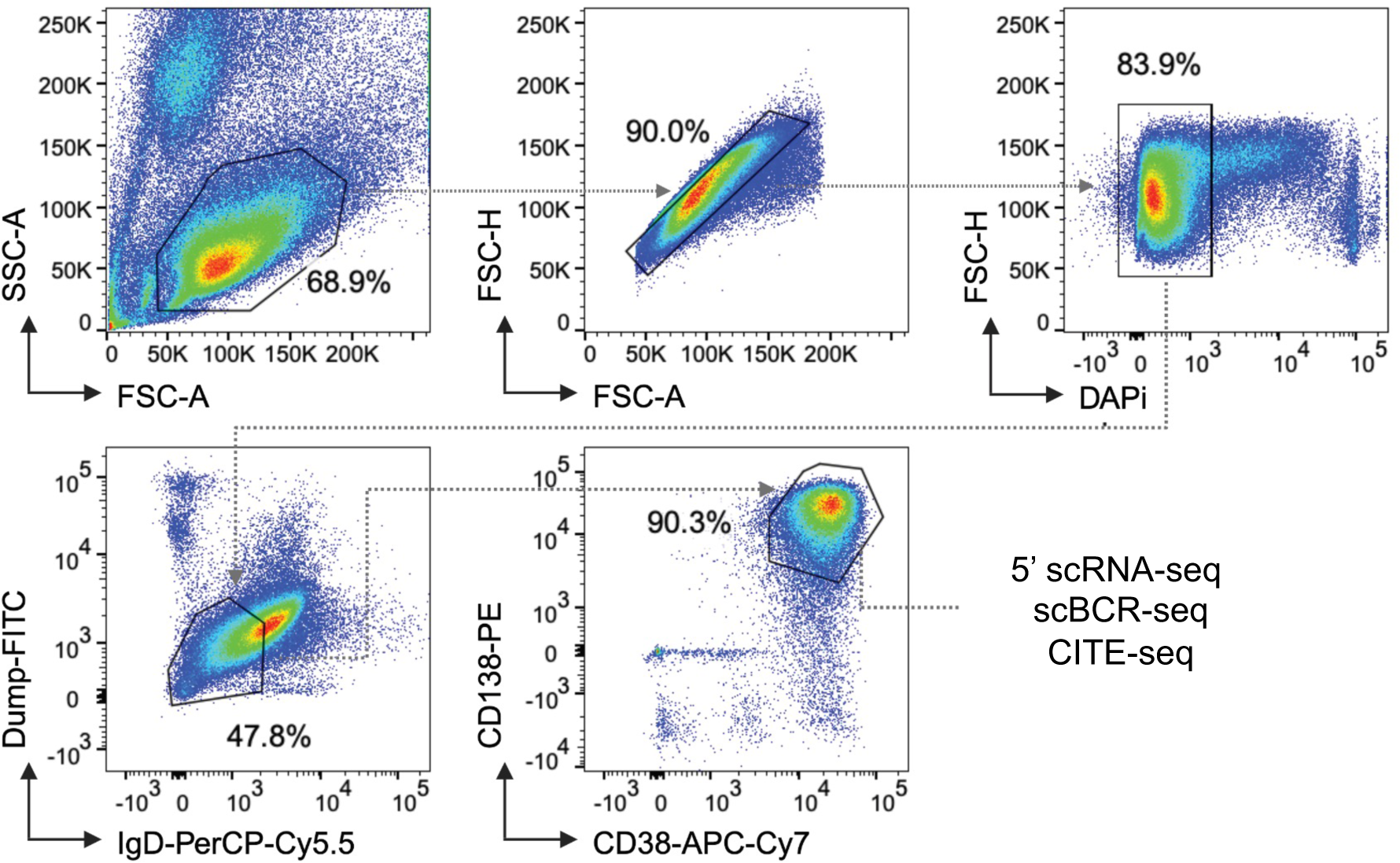
Gating strategy. FACS sorting of magnetically enriched CD138^+^ human bone marrow plasma cells. Dump channel contains anti-CD3 and anti-CD14 antibodies.

**Figure S2.**
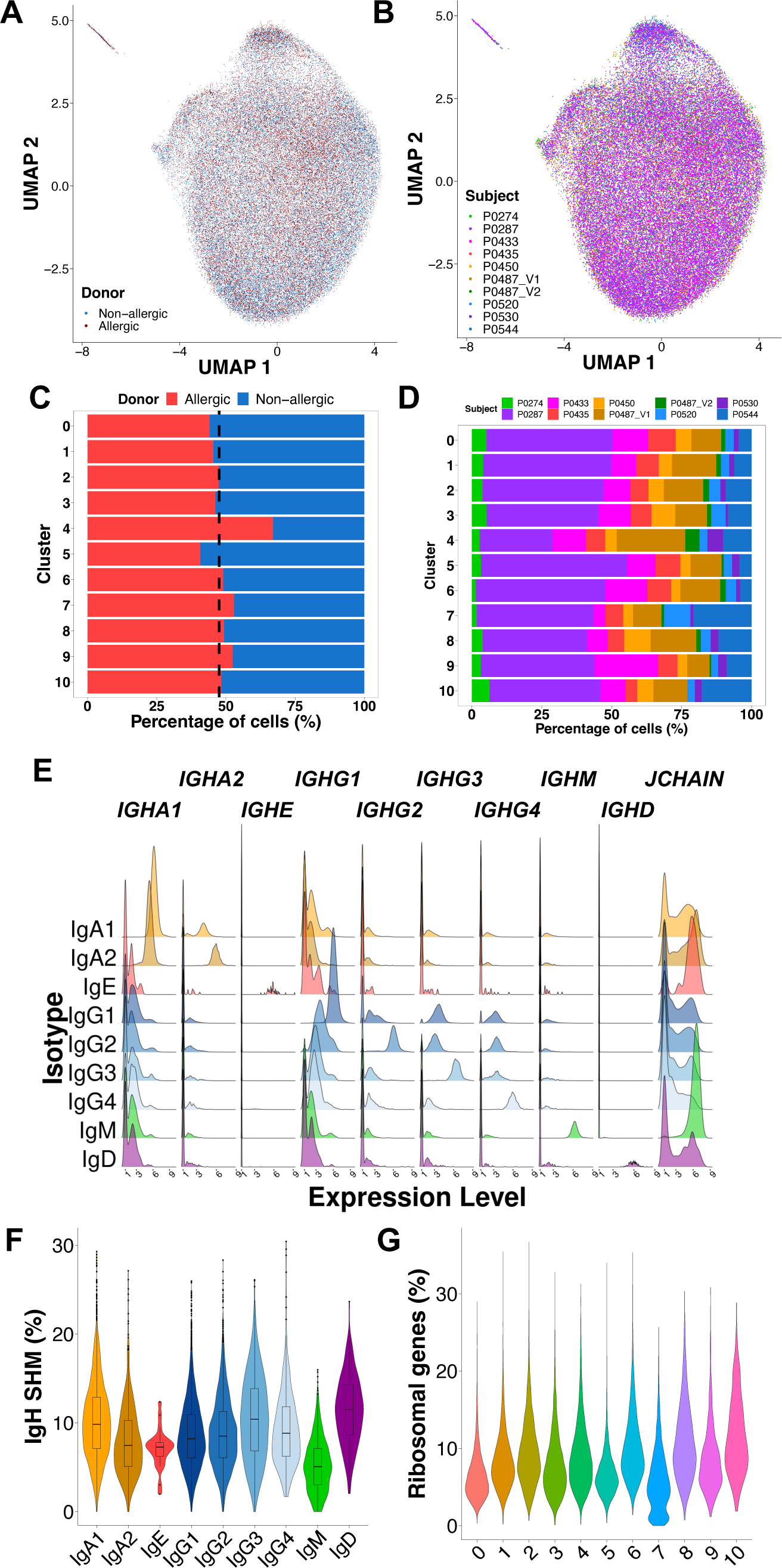
Quality control of clustered cells. **(A)** UMAP projection of BMPCs, colored by donor’s allergy status. **(B)** Proportions of cells in each cluster by donor’s allergy status. Black dashed line represents expected proportion based on input number of cells per allergy status (n = 34,991 for allergic, n = 38,569 for non-allergic). **(C)** UMAP projection of BMPCs, colored by donor. **(D)** Proportions of cells in each cluster by donor. **(E)** Ridge plots for IgH gene expression on BMPCs of different IgH isotypes. **(F)** Heavy chain somatic hypermutation (IgH SHM) percentage of BMPCs of different IgH isotypes. **(G)** Percentage of ribosomal genes in the transcriptome of each cluster.

**Figure S3.**
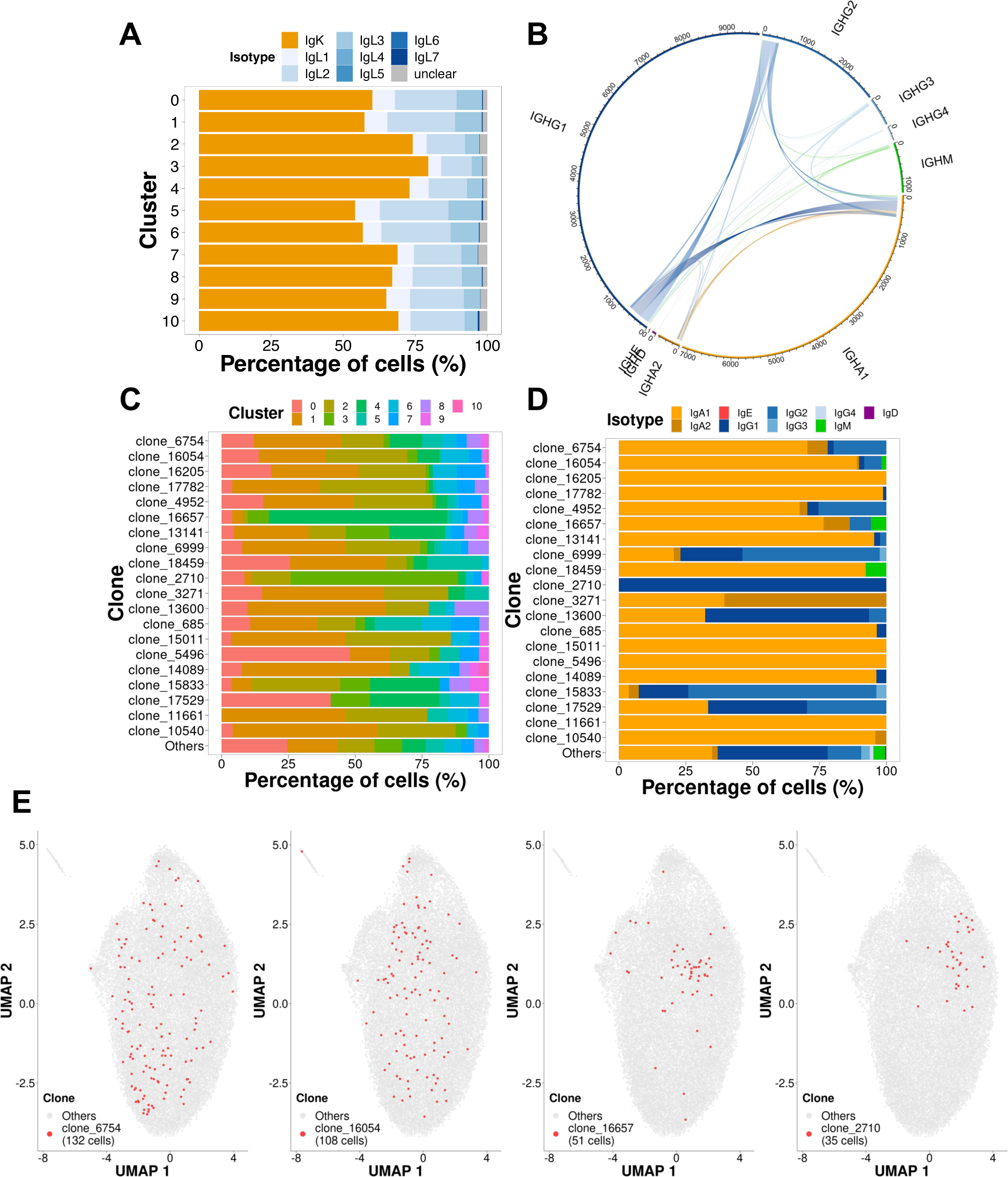
**(A)** Proportion of cells in each cluster by IgL isotype. **(B)** Clonal relationships between BMPCs of different isotypes. Lines connect cells that have the same IgH V and J segments, and over 82% identity in CDRH3 (Hamming distance determined by Shazam = 0.18). Line end width is proportional to the number of cells in the clonal family. **(C-D)** Top 20 most expanded clonal families. Composition of each family is colored by cluster identity (C) and IgH isotype (D). Remaining aggregated BMPCs are included for reference (“Others”). **(E)** UMAP projection of clustered BMPCs. Largest clone families (6754 & 16054) and predominantly mono-isotypic clone families (16657 & 2710) showed in **(C)** and **(D)** are highlighted in red.

**Figure S4.**
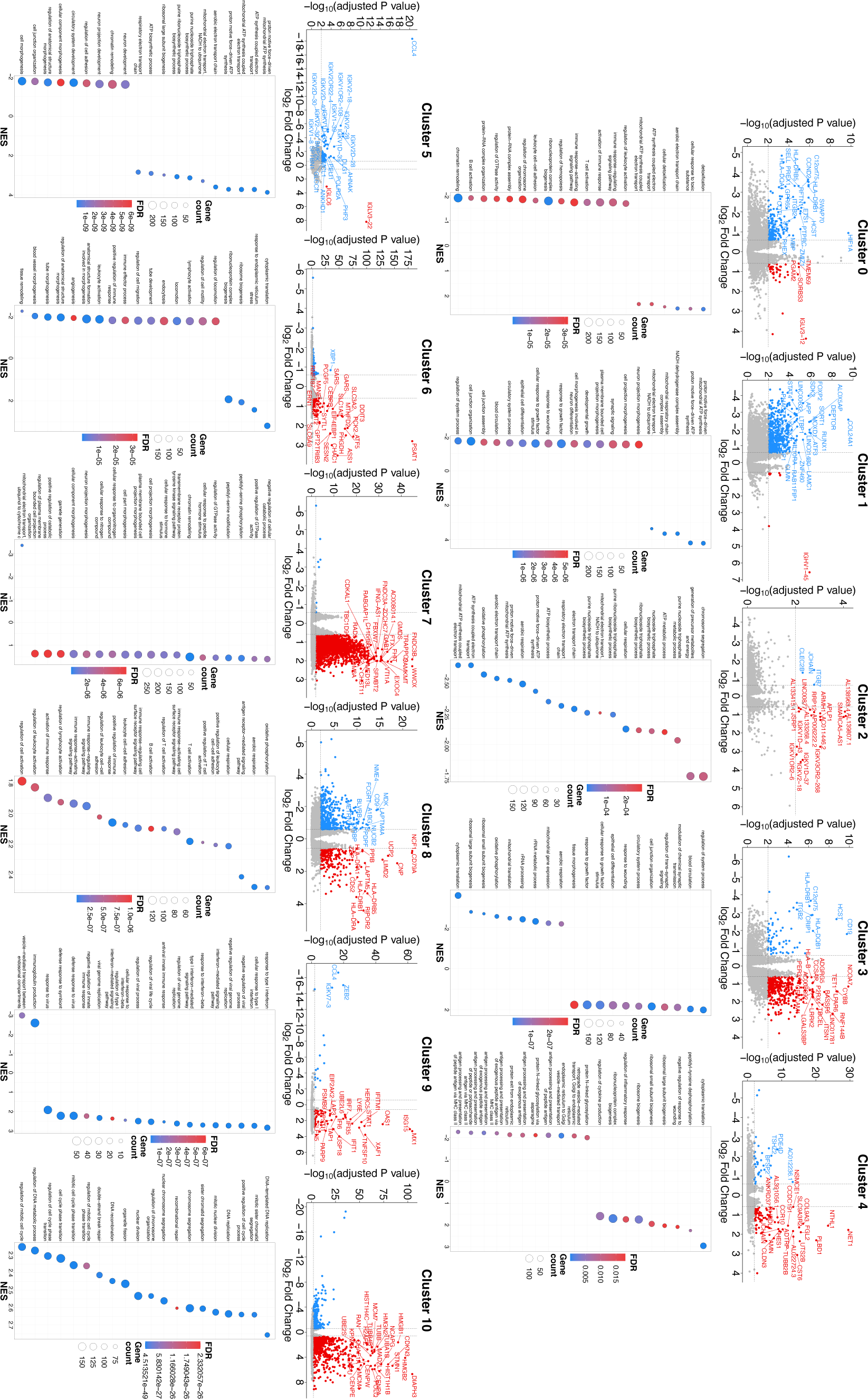
Pathway analysis for clusters of BMPCs. Differential expression gene testing was performed on BMPCs pseudobulked by donor and cluster identity. For each set of genes, gene set enrichment analysis was performed. Volcano plots and the top 20 GSEA results by false discovery rate (FDR) are shown for each cluster.

**Figure S5.**
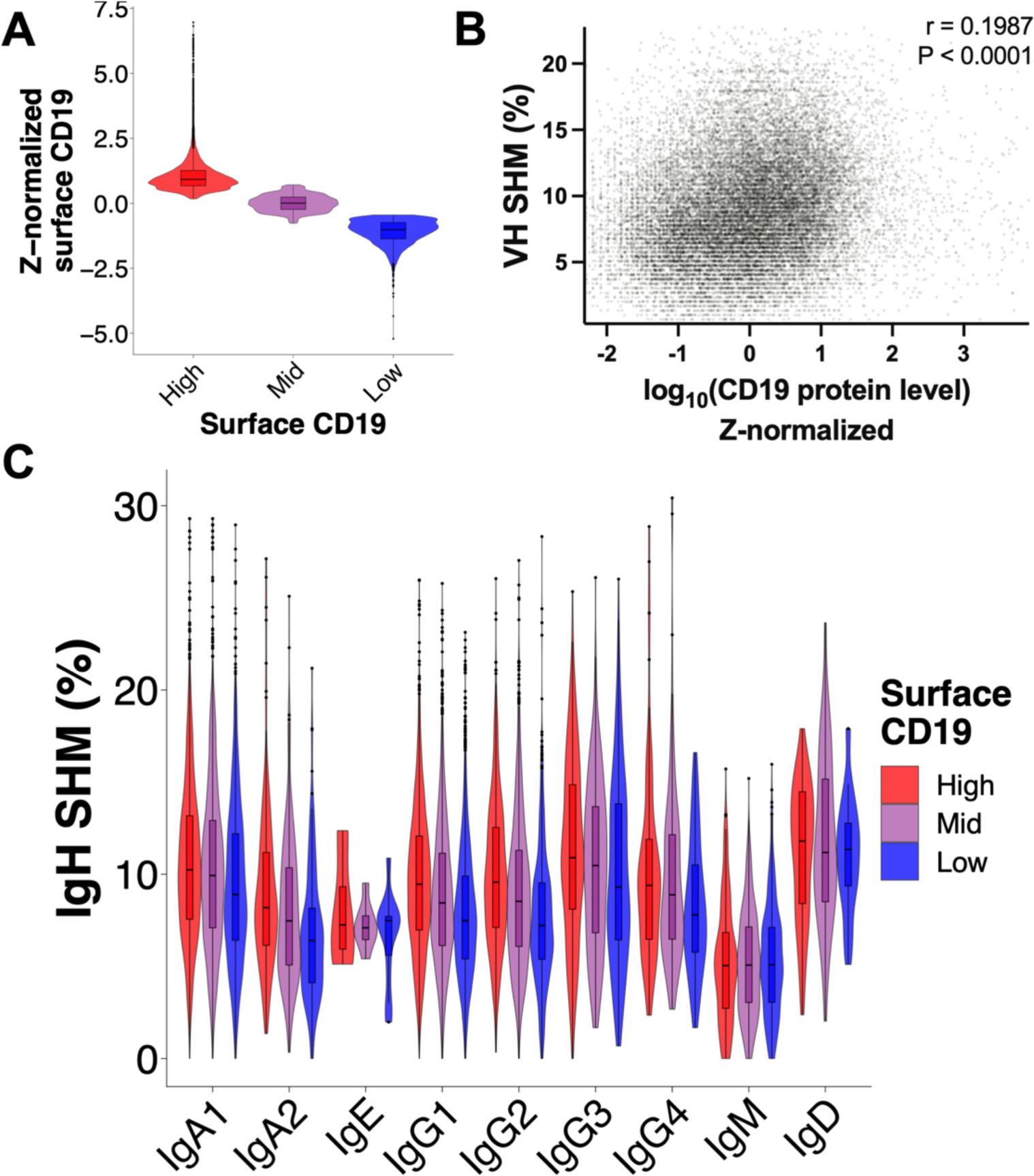
Surface CD19 expression and SHM burden are correlated. **(A)** Normalized surface CD19 expression on CD19 expression categories among BMPCs. **(B)** Correlation between normalized surface CD19 expression and IgH SHM (n = 23,289). Spearman correlation was performed. 99.5% of the distribution in the X axis and 99.5% of the distribution in the Y axis are shown. **(C)** IgH SHM of BMPCs of different isotypes, subdivided by surface CD19 expression categories. Sample sizes for CD19 high/mid/low are: IgA1 = 3,137 / 3,497 / 1,892; IgA2 = 165 / 218 / 141; IgE = 4 / 8 / 10; IgG1 = 2,137 / 3,411 / 3,401; IgG2 = 926 / 1,188 / 760; IgG3 = 160 / 295 / 230; IgG4 = 103 / 128 / 73; IgM = 252 / 485 / 457; IgD = 42 / 41 /30).

**Figure S6.**
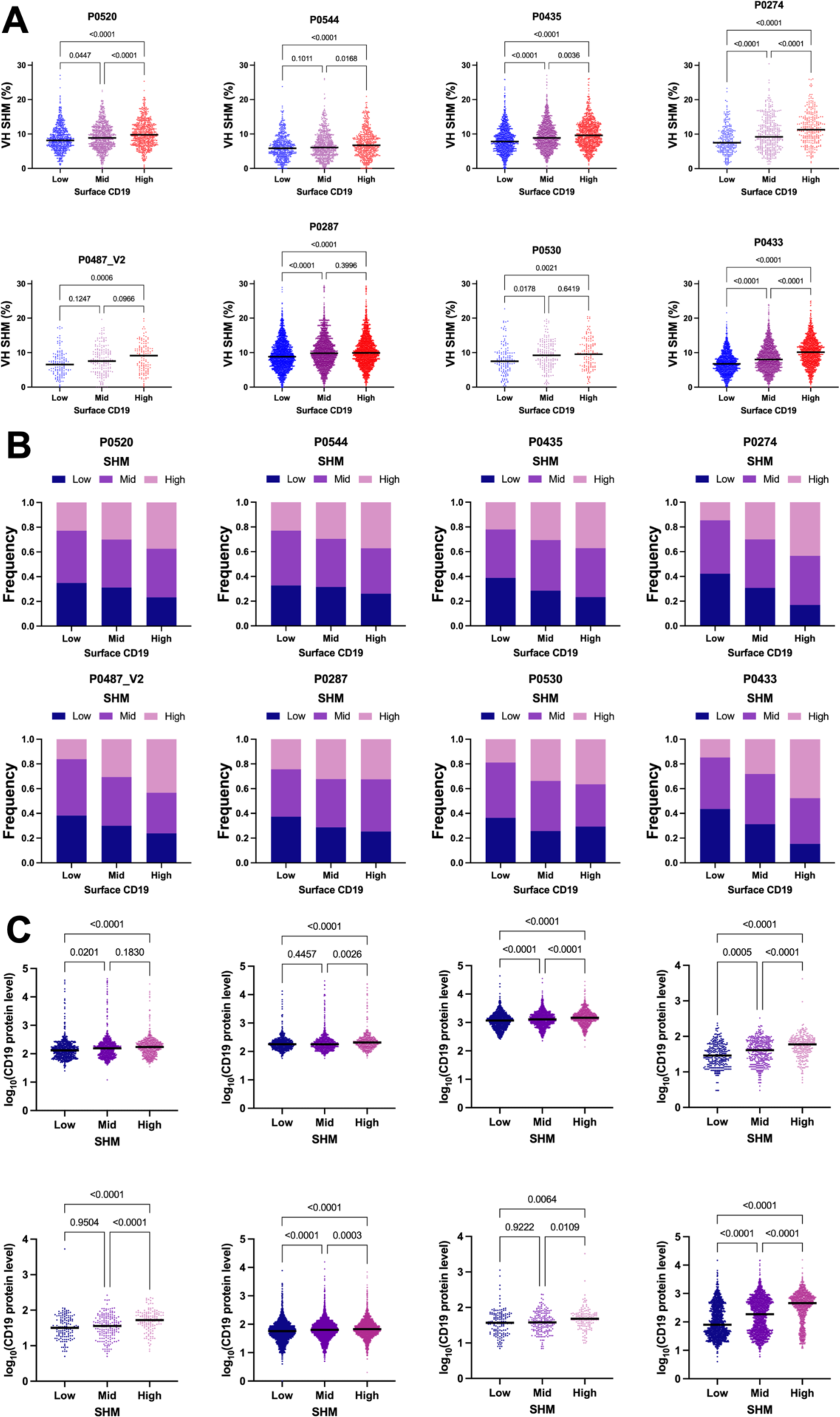
Features indicative of BMPC origin are retained at the donor level. **(A)** IgH SHM per surface CD19 expression bins per donor. Kruskal-Wallis test was used to assess statistical significance. **(B)** Frequency of IgH SHM bins per cluster per donor. **(C)** Surface CD19 expression per IgH SHM bins per donor. Kruskal-Wallis test was used to assess statistical significance.

**Figure S7.**
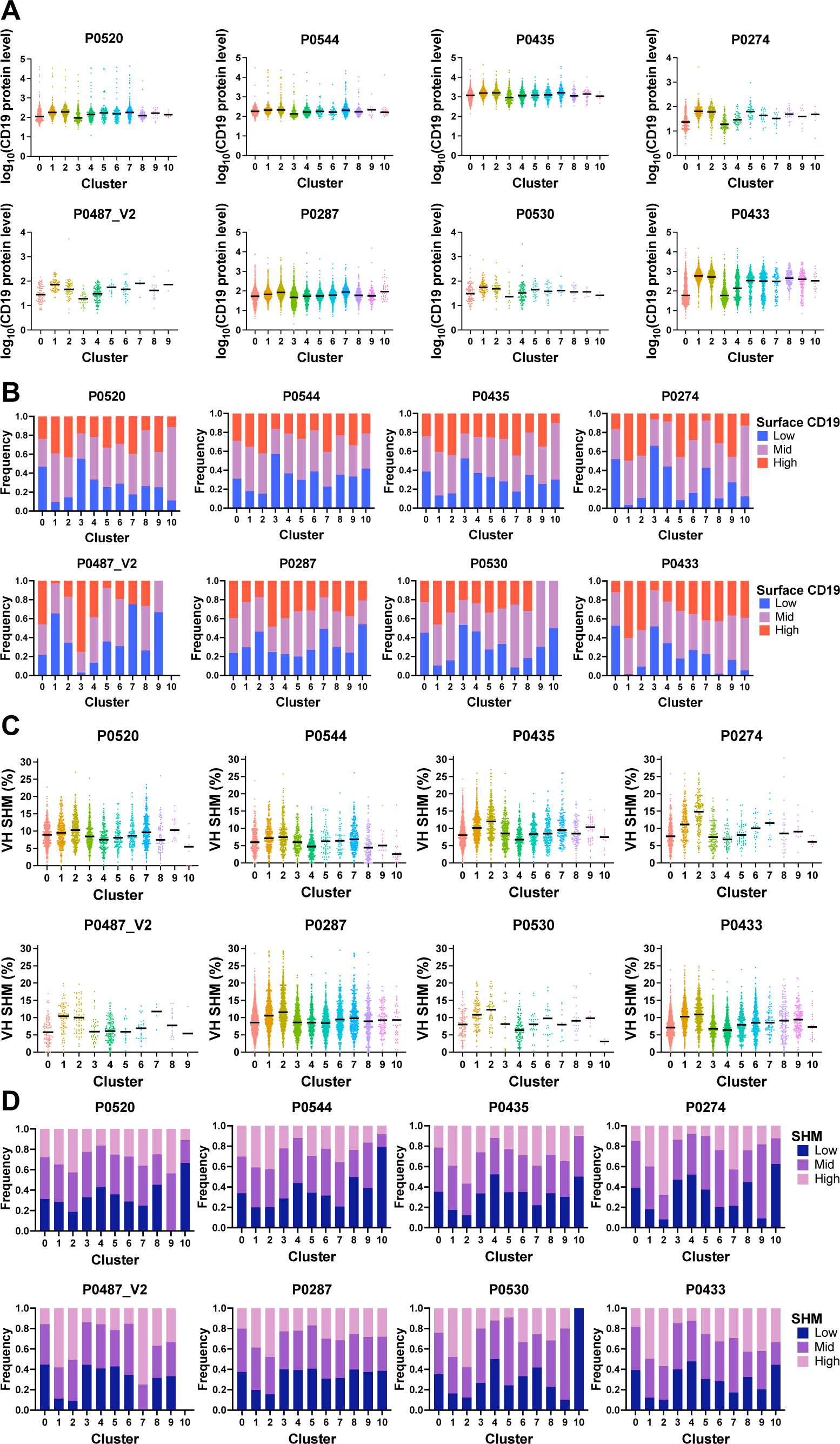
Surface CD19 expression and SHM levels are retained at the donor level. **(A)** Surface CD19 expression on BMPCs per cluster per donor. **(B)** Frequency of surface CD19 expression bins per cluster per donor. **(C)** Somatic hypermutation (SHM) in IgH chain per cluster per donor. **(D)** Frequency of IgH SHM bins per surface CD19 expression bins per donor.

**Figure S8.**
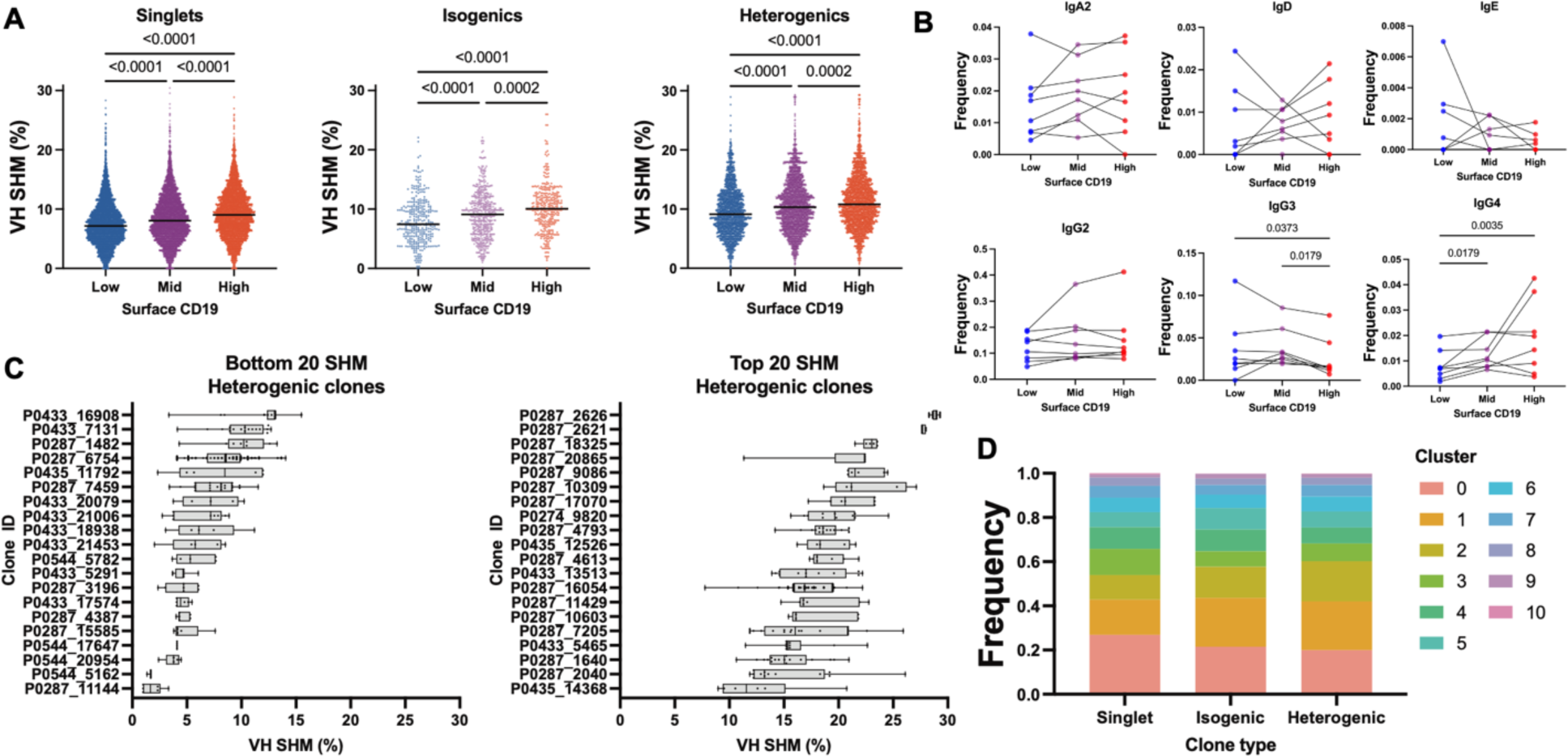
Features of clonal structure related to BMPC origins. **(A)** IgH somatic hypermutation (SHM) per surface CD19 bin in clonal families with different clonal structures. Kruskal-Wallis test was used to assess statistical significance. **(B)** Frequency of surface CD19 bins among IgA2, IgD, IgE, and IgG2/3/4 BMPCs. Lines connecting dots represent an individual donor. Friedman test was used to assess statistical significance. Only statistically significant differences are shown. **(C)** IgH SHM in the 20 least mutated and the 20 most mutated heterogenic clones (related to Fig. 3I). **(D)** Frequency of cluster identities among clonal families with different clonal structures.

**Figure S9.**
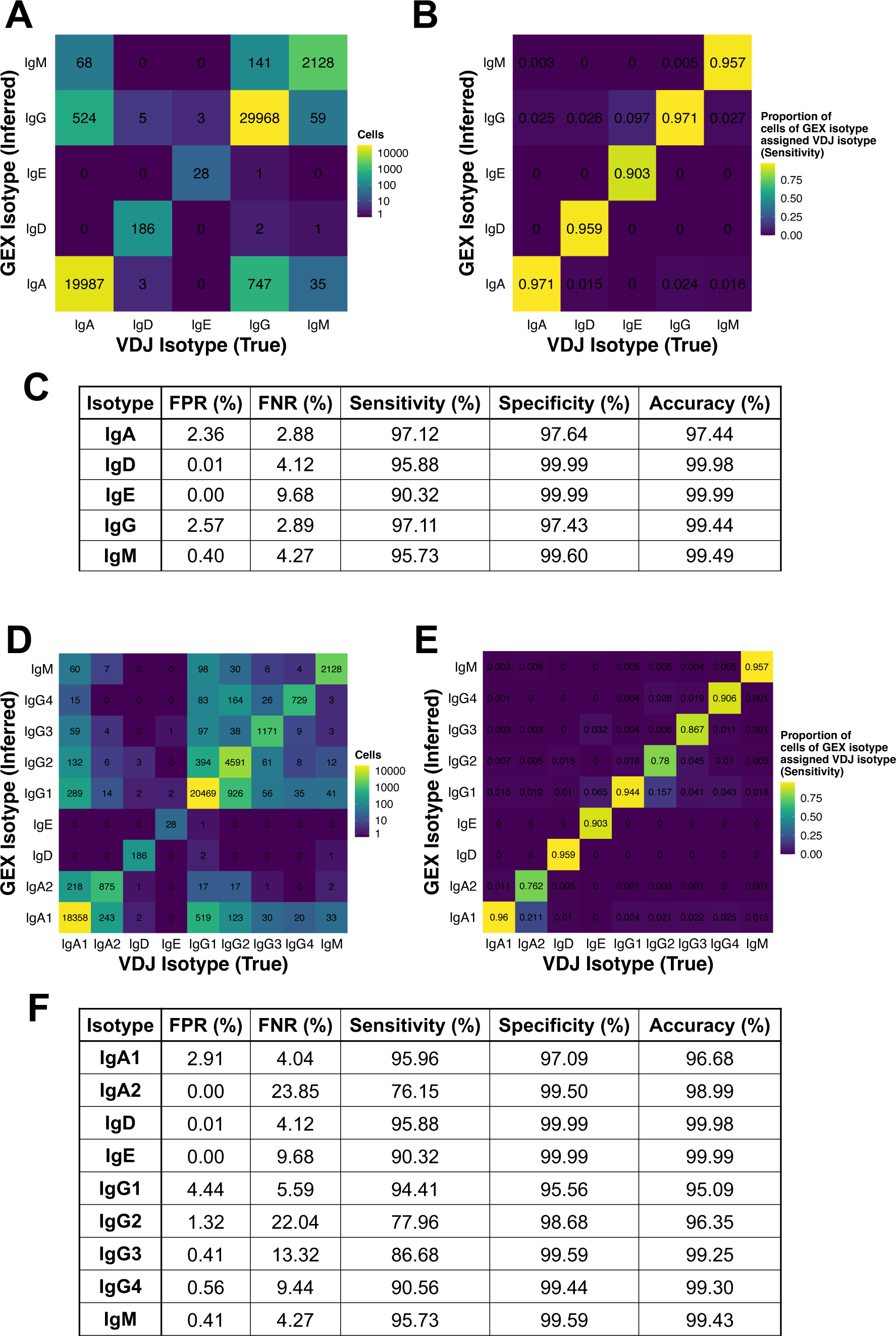
Performance of GEX-based isotype calling when benchmarked against VDJ-based isotype calling. **(A)** Number of cells assigned IgH isotype class based on gene expression (GEX) library, compared to “true” isotype class calling based on VDJ library (n = 49,667). **(B)** Percentage of cells of a given IgH isotype class (as determined by GEX library) that are inferred to pertain to each IgH isotype class based on VDJ library (n = 49,667). **(C)** Performance metrics of GEX-based isotype calling when benchmarked against VDJ library callings at the IgH class level. FPR = False positive rate. FNR = False negative rate. **(D)** Number of cells assigned IgH isotype subclass based on GEX library, compared to “true” isotype subclass calling based on VDJ library (n = 49,667). **(E)** Percentage of cells of a given IgH isotype subclass (as determined by GEX library) that are inferred to pertain to each IgH isotype subclass based on VDJ library (n = 49,667). **(F)** Performance metrics of GEX-based isotype calling when benchmarked against VDJ library callings at the IgH subclass level. FPR = False positive rate. FNR = False negative rate.

**Fig. S10.**
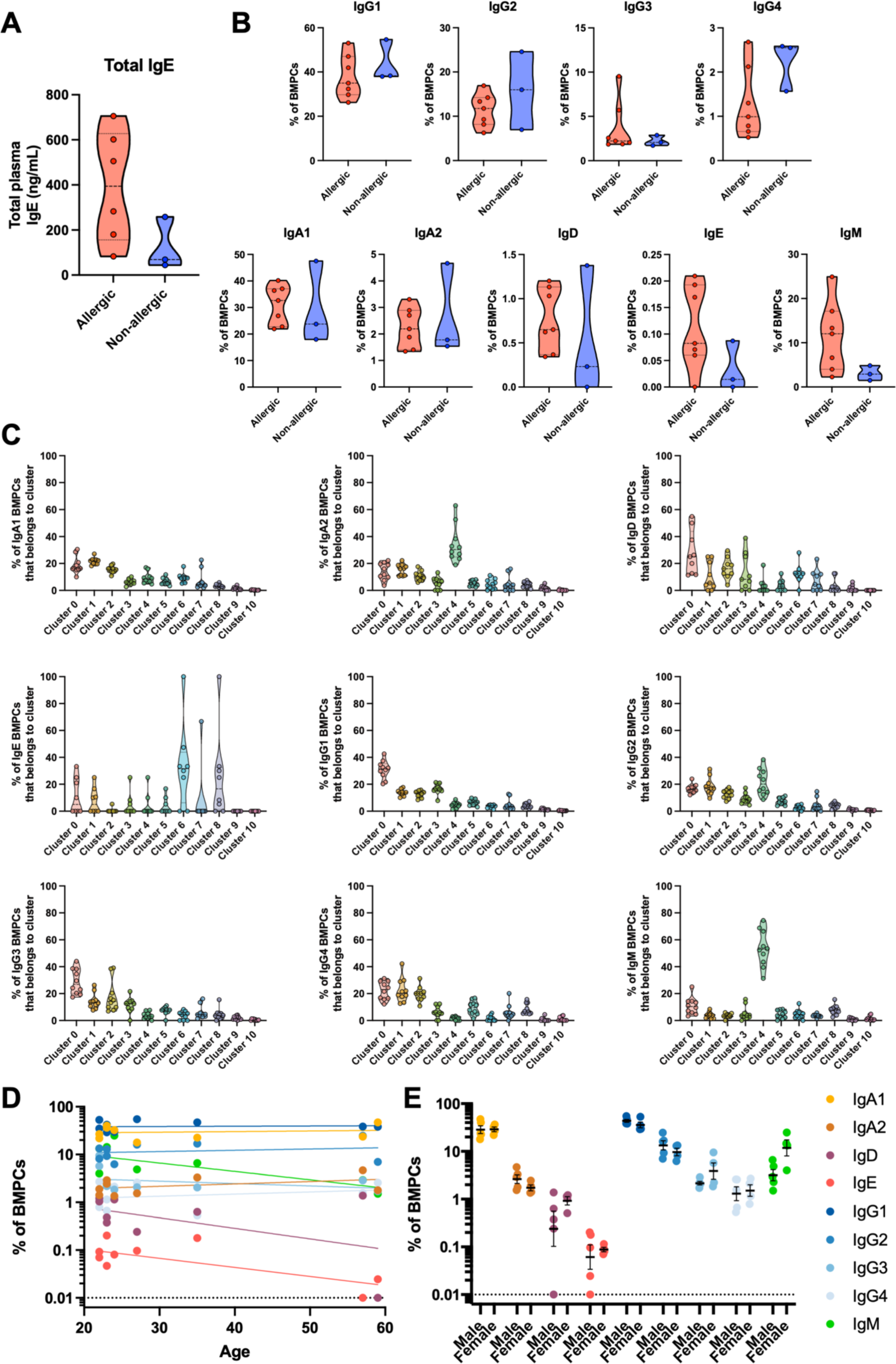
Plasma cell heavy chain isotype proportions in bone marrow. **(A)** Total IgE concentrations in plasma of donors. **(B)** Frequency of BMPCs of different IgH isotypes among donors. **(C)** Frequency of BMPCs of different IgH isotypes among clusters. **(D)** Age is not a covariate that explains differences in BMPC isotype proportions among allergic and non-allergic individuals. Spearman correlation was performed. **(E)** Sex is not a covariate that explains differences in BMPC isotype proportions among allergic and non-allergic individuals. Paired Friedman test was performed to identify differences.

**Figure S11.**
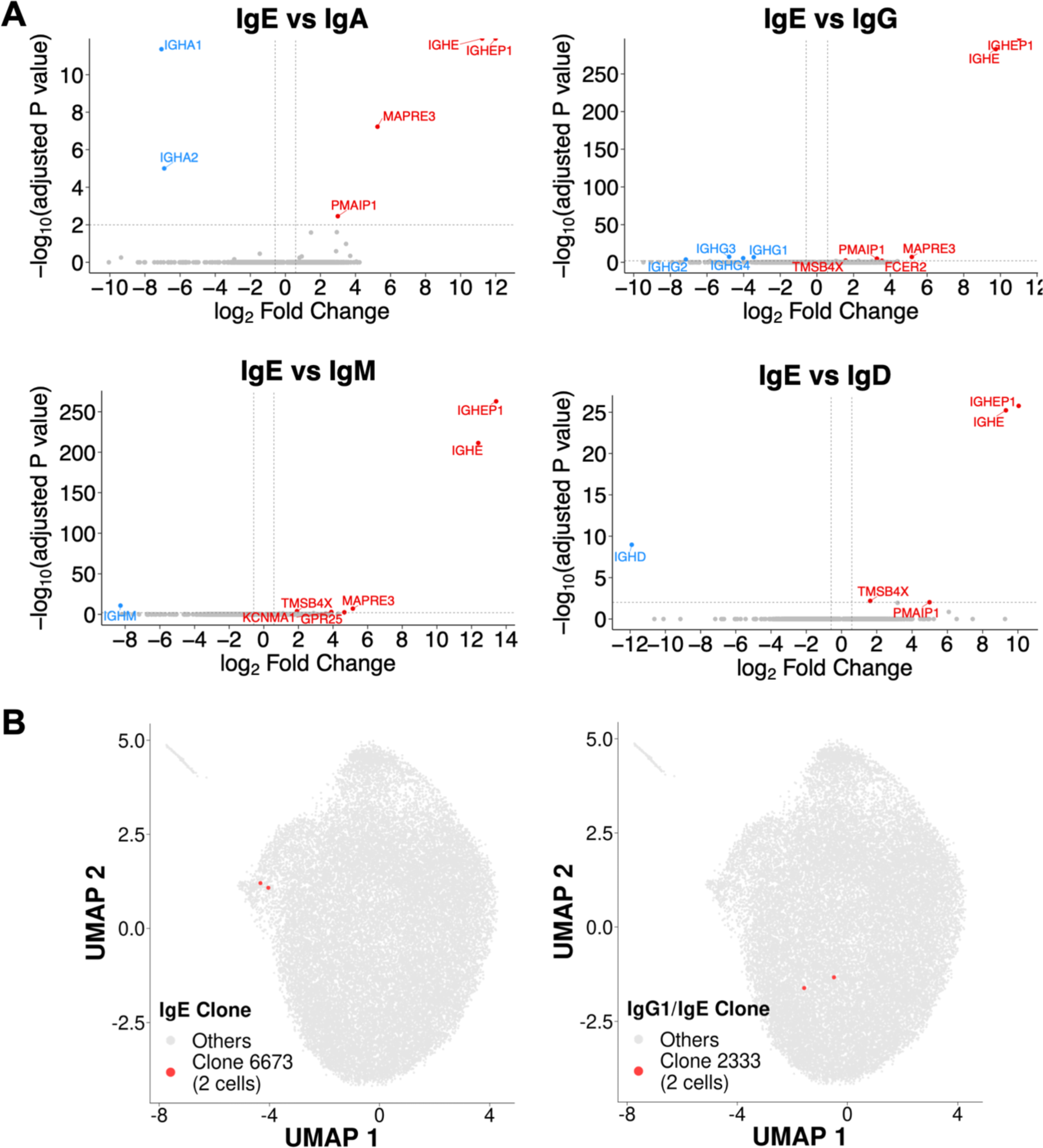
IgE BMPC DEGs and clonal families. **(A)** Volcano plots of differentially expressed genes among IgE cells and different isotype classes. **(B)** UMAP projection of clustered BMPCs, highlighting clones that include at least 1 IgE BMPC.

**Figure S12.**
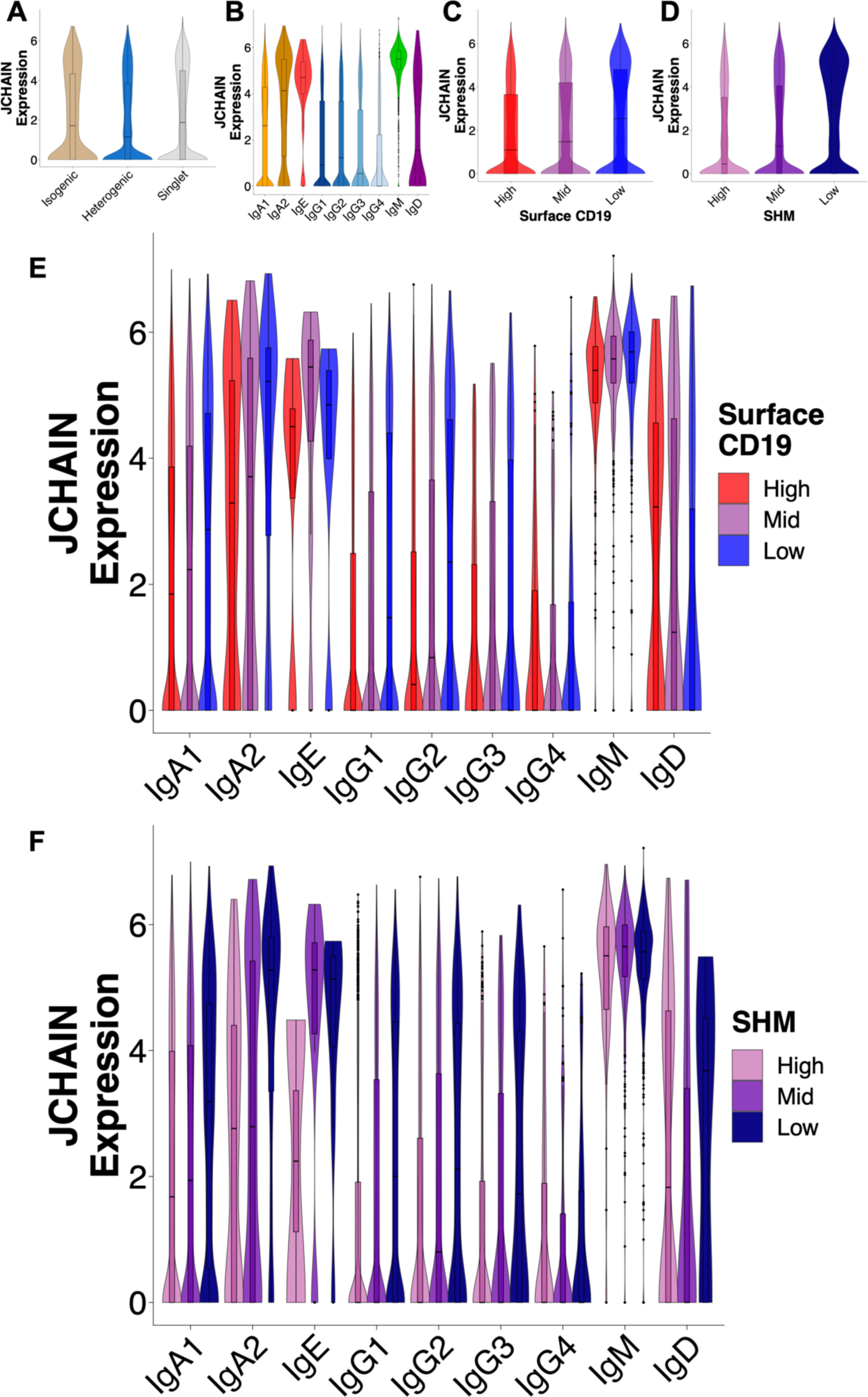
J chain expression among BMPCs. **(A)** Expression of *JCHAIN* among clonotypes of different structures. **(B)** Expression of *JCHAIN* among BMPCs of different IgH subclasses. **(C)** Expression of *JCHAIN* among BMPCs belonging to different surface CD19 expression categories. **(D)** Expression of *JCHAIN* among BMPCs belonging to different IgH SHM categories. **(E)** *JCHAIN* expression among BMPCs of different isotypes, subdivided by surface CD19 expression categories. Sample sizes for CD19 high/mid/low are: IgA1 = 3,137 / 3,497 / 1,892; IgA2 = 165 / 218 / 141; IgE = 4 / 8 / 10; IgG1 = 2,137 / 3,411 / 3,401; IgG2 = 926 / 1,188 / 760; IgG3 = 160 / 295 / 230; IgG4 = 103 / 128 / 73; IgM = 252 / 485 / 457; IgD = 42 / 41 /30). **(F)** *JCHAIN* expression among BMPCs of different isotypes, subdivided by IgH SHM categories. Sample sizes for IgH SHM high/mid/low are: nIgA1 = 3,246 / 3,341 / 1,939; nIgA2 = 107 / 204 / 213; IgE = 2 / 11 / 9; IgG1 = 2,222 / 3,725 / 3,002; IgG2 = 804 / 1,213 / 857; IgG3 = 314 / 109 / 96; IgG4 = 99 / 109 / 96; IgM = 74 / 396 / 724; IgD = 72 / 30 / 11).

**Table S1.** Cohort demographics.

| Subject ID | Sex | Age (years) | Allergy status | Food allergies | Aeroallergy | Other allergies or intolerances | Reaction type / Time since last reaction | Total IgE (ng/mL) | Peanut-specific IgE* (kUA/L) |
| --- | --- | --- | --- | --- | --- | --- | --- | --- | --- |
| P0274 | M | 57 | Non-allergic | None | No | No | n/a | 258.21 | n/a |
| P0287 | M | 59 | Non-allergic | None | No | No | n/a | 68.86 | n/a |
| P0450<sup>†</sup> | M | 27 | Non-allergic | None | No | No | n/a | 44.03 | n/a |
| P0435 | M | 23 | Allergic | Peanuts, tree nuts, sesame | Yes | No | Local, no history of anaphylaxis | 705.92 | n/a |
| P0487 | F | 23 | Allergic | Peanuts, sesame, egg, dairy | No | Penicillins | Systemic / n.r. | 505.59 | 69.5 |
| P0530 | F | 24 | Allergic | Peanuts, tree nuts, egg, dairy, mussels, oysters, clams | Yes | No | Systemic / 22 months | 180.37 | 0.17 |
| P0433 | M | 35 | Allergic | Peanuts | Yes | No | Systemic / >10 years | 283.29 | 0.37 |
| P0520<sup>‡</sup> | F | 22 | Allergic | Berries, tomatoes | Yes | Latex | n.r. | 82.95 | 0.17 |
| P0544<sup>‡</sup> | F | 22 | Allergic | Peanuts | Yes | Penicillin, mosquitoes | Systemic / >2 years | 601.34 | >100 |
| P0069<sup>‡§</sup> | F | 34 | Allergic | Peanuts, legumes, soy | No | Halothane, nickel | Systemic / 2 months | 64.57 | 32.5 |
| P0263<sup>‡§</sup> | F | 28 | Allergic | Dairy | No | No | n.r. | 523.26 | n/a |

\* Measured by ImmunoCAP.
<sup>†</sup> Populations FACS-sorted by CD19 surface expression before sequencing. Data on CD19 expression not used for analyses.
<sup>‡</sup> VDJ rearrangements cloned into IgG1 for peanut binding testing.
<sup>§</sup> Data not included in clustering because of strong batch effects.
n/a: Not applicable
n.r.: Not reported

